# Accurate Prediction of ecDNA in Interphase Cancer Cells using Deep Neural Networks

**DOI:** 10.1101/2025.06.23.661188

**Authors:** Utkrisht Rajkumar, Gino Prasad, Ellis J Curtis, Ivy Tsz-Lo Wong, Xiaowei Yan, Shu Zhang, Lotte Brückner, Kristen Turner, Julie Wiese, Justin Wahl, Sihan Wu, Jessica Theissen, Matthias Fischer, Howard Y. Chang, Anton G. Henssen, Paul S. Mischel, Vineet Bafna

## Abstract

Oncogene amplification is a key driver of cancer pathogenesis and is often mediated by extrachromosomal DNA (ecDNA). EcDNA amplifications are associated with increased pathogenicity of cancer and poorer outcomes for patients. EcDNA can be detected accurately using fluorescence in situ hybridization (FISH) when cells are arrested in metaphase. However, the majority of cancer cells are non-mitotic and must be analyzed in interphase, where it is difficult to discern extrachromosomal amplifications from chromosomal amplifications. Thus, there is a need for methods that accurately predict oncogene amplification status from interphase cells.

Here, we present interSeg, a deep learning-based tool to cytogenetically determine the amplification status as EC-amp, HSR-amp, or not amplified from interphase FISH images. We trained and validated interSeg on 652 images (40,446 nuclei). Tests on 215 cultured cell and tissue model images (9,733 nuclei) showed 89% and 97% accuracy at the nuclear and sample levels, respectively. The neuroblastoma patient tissue hold-out set (67 samples and 1,937 nuclei) also revealed 97% accuracy at the sample level in detecting the presence of focal amplification. In experimentally and computationally mixed images, interSeg accurately predicted the level of heterogeneity. The results showcase interSeg as an important method for analyzing oncogene amplifications.

## INTRODUCTION

Oncogene amplification is a key driver of cancer pathogenesis^1^. Focal oncogene amplifications can occur within specific chromosomes as homogeneously staining regions^2,3^ (HSR) or as extrachromosomal DNA^4^ (ecDNA), which are circular, acentric molecules that replicate independently and segregate randomly in daughter cells^5^. EcDNAs are present in a third of all samples, and in two-thirds of cancer subtypes^6^. They are especially frequent in glioblastoma^7^, neuroblastoma^8^, and esophageal carcinoma^4,6^, but have also been detected in pre-cancerous lesions^9^. Compared to other intrachromosomal focal amplifications, ecDNAs are associated with increased pathogenicity of cancer and poorer outcomes for patients^6^. Thus, there is an important need for methods and tools to detect focal amplifications in tumor cells and classify their location as being intrachromosomal or extrachromosomal.

Sequence-based methods^10,11^ analyze the patterns of genomic reads sampled from a tumor genome and mapped to a normal reference to (a) identify copy number patterns indicative of focal amplification, (b) use focally amplified regions as seeds, and (c) utilize discordantly paired-reads to explore the fine genomic structure of focal amplifications. The presence of discordant reads that represent a cyclic structure are highly indicative of ecDNA structure^6^. Sequence-based methods can also reliably distinguish ecDNA from stable chromosomal amplifications (displaying as HSRs) formed by breakage fusion bridge cycles and other mechanisms^3^. However, HSRs may also be formed when ecDNA re-integrate into chromosomes in response to the cellular environment^12^. The HSRs formed by re-integrated ecDNA retain their sequence features, making it difficult for sequence-based methods to predict the amplification mechanism.

Fluorescent and DAPI imaging of DNA in metaphase spreads are currently the gold-standard for determining the location (intra- or extrachromosomal) of focal amplification. EcDNAs appear as hundreds of tiny faint DNA particles, detached from the compacted chromosomes seen in metaphase. Fluorescently labeled DNA FISH probes for specific genes can additionally determine if the ecDNAs carry those genes. A deep-learning method, ecSeg, was successfully utilized to semantically segment images of metaphase cells and annotate the pixels representing ecDNA^13^. However, capturing cells in metaphase requires synchronization of cells, which is typically possible only in cultured cell lines. In clinical practice, cells are harvested from patient tumor tissue and readily archived as flash-frozen tissue sample, or as formalin-fixed paraffin-embedded samples. The majority of cells are non-mitotic and must be analyzed in interphase, where the DNA is loosely arranged inside an intact nuclear membrane. This makes it extremely challenging to identify ecDNA, even for a trained eye.

In this work, we discern HSR and ecDNA amplifications using the unique fluorescent staining patterns of amplicons in interphase nuclei. We present interSeg, a deep learning-based tool to cytogenetically determine amplification status of a target FISH probe. interSeg relies on two independent deep learning modules: ecSeg-c and ecSeg-i. EcSeg-c uses centromeric and target FISH probes to determine if the target is amplified. The additional centromeric probe provides a control for aneuploidy, whole genome duplication and overlapping cells, which may result in higher number of FISH foci. EcSeg-i determines the mode of amplification as ecDNA or HSR, assuming focal amplification of the target, and requires only the target FISH probe.

## RESULTS

We modeled ecDNA detection in interphase nuclei as a problem of nuclear classification, where each interphase nucleus was assigned to one of three categories (Figure 1a): amplification on ecDNA (‘EC-amp’); intrachromosomal amplification, described by a homogeneously stained region (‘HSR-amp’); or no amplification of the target probe (‘no-amp’). Each cytogenetic image itself contained a collection of interphase nuclei and were found to have different characteristics depending on the source. Therefore, we first collected representative cytogenetic images from different sources to create a dataset.

**Figure 1:**
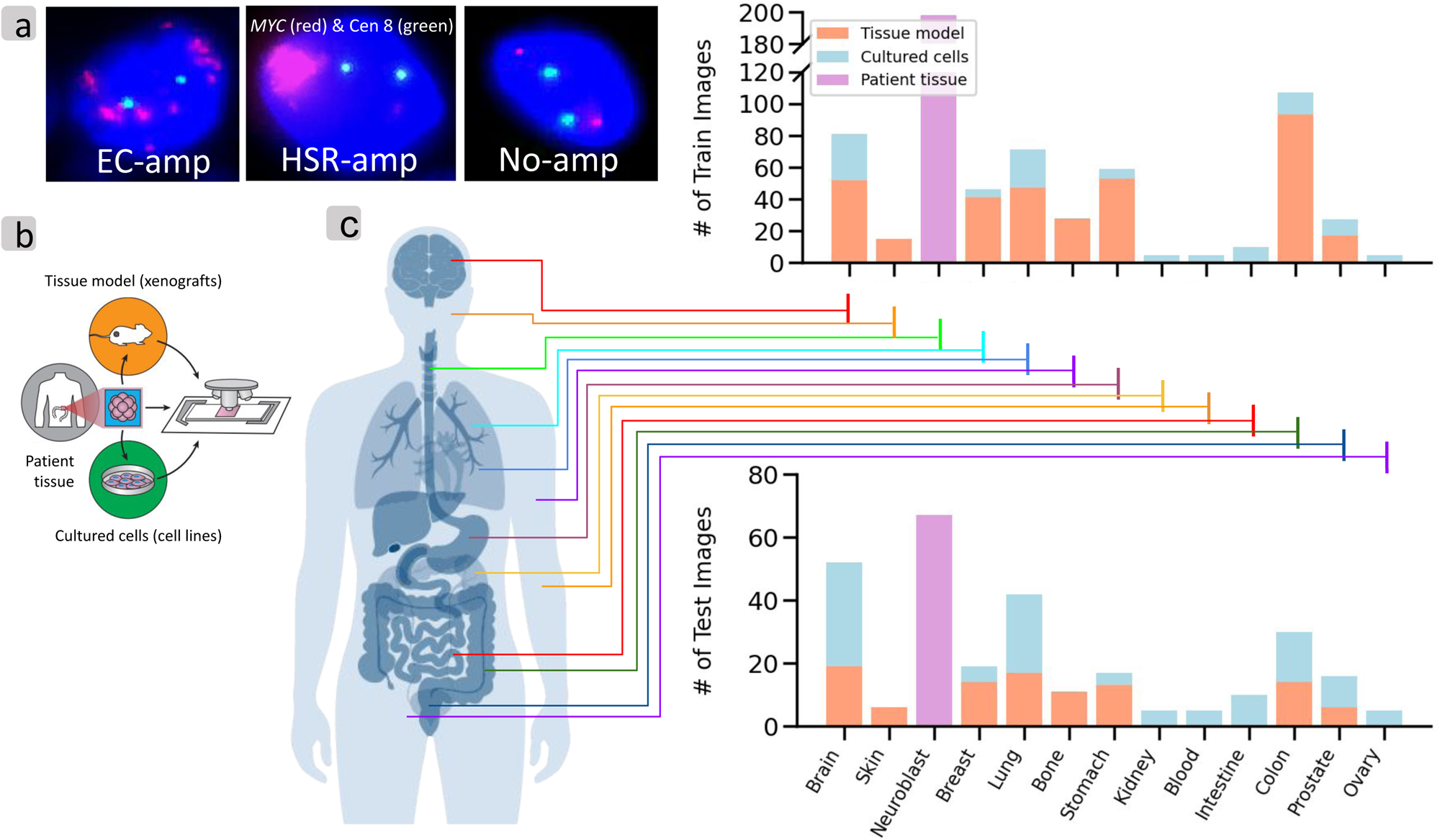
Data overview and Analysis pipeline. (a) Examples of interphase cells with EC-amp, HSR-amp, No-amp (*MYC* red, Centromere 8 green). (b) Image acquisition protocols for various tissue modalities. (c) Distribution of tissue types and image acquisition protocols in analyzed cell lines and patient tissues, for the training and hold-out test set.

### Dataset overview

We obtained images from 13 different tissue types using three different protocols (Figure 1b). ‘Cultured cells’ refer to cells grown outside of their natural environment, typically in a culture, and plated on a slide prior to image acquisition. This results in images with dissociated and sparse nuclei. ‘Tissue models’ are cells obtained from xenografts, which are tissues transplanted from human to mouse, then biopsied for image acquisition. This results in images with more tightly packed cells. ‘Patient tissue’ corresponds to tightly packed cells in tissue sections from tumor biopsies (Supplementary Figure 1 and Methods), but may have more heterogeneity and a higher fraction of normal tissue cells. We obtained 231 cultured cell images and 443 filtered tissue model images from 32 unique cell lines, and 265 filtered images from patient derived (see Methods) neuroblastoma (NB) samples. In addition, we utilized 60 ‘mixed’ images (765 nuclei) from a special tagging experiment designed to test performance in heterogeneous samples containing both ecDNA and HSR (See Methods).

We used whole genome sequencing (WGS) to identify the amplified oncogene in the cultured cell and tissue model cell lines, as described in earlier publications^6^. We then probed for these amplified genes using FISH probes in DAPI stained metaphase spreads where the chromosomes are compacted, and the nuclear location of the FISH probe can be unambiguously determined. This provided the truth set for whether the oncogene was amplified on ecDNA or HSR^6^. A few cell lines were probed for more than one oncogene (e.g. H716 for *FGFR2* and *MYC*) to obtain 39 unique cell line-oncogene pairs. The cell line-oncogene pairs (𝑙, 𝑔) from the cultured cell lines and tissue-models were assigned a label 𝐿(𝑙, 𝑔) as being one of EC-amp, HSR-amp, or no-amp. Correspondingly, each nucleus with a fluorescent label for gene 𝑔 in an interphase image of cell line 𝑙 also received the label 𝐿(𝑙, 𝑔), providing us with a data set of nuclei that could be utilized for training, validation, and testing. For example, we labeled all nuclei in the COLO320HSR cell line as HSR-amp for the oncogene *MYC*.

We trained ecSeg-i and ecSeg-c separately. For ecSeg-i, we performed a 50-50 training/validation hold-out set split on the 231 cultured cell images, and a 75-25 training/validation hold-out set split on the 443 tissue model images. In total, we utilized 459 images (35,096 nuclei) for training/validation, and 215 images for hold-out testing (Supplementary Table 1). Finally, we utilized the NB patient tissue images as analysis sets for biological interpretation (265 images, 7,466 nuclei). Notably, the trained neural networks never accessed the 215 hold-out test images from cultured cells and tissue models, 60 mixed images, or the 265 patient tissue images during training or validation of ecSeg-i.

EcSeg-c requires centromeric and target FISH probes, and it returns a classification of ‘focal-amp’ or ‘no-focal-amp’ for each nucleus. 392 of the 443 tissue model images had a centromeric probe and met our centromeric quality score criterion. We labeled images with HSR-amp and EC-amp classifications as ‘focal-amp’ for ecSeg-c training. Other images were labeled ‘no-focal-amp’. Of the 392 images, 95 images (7,501 nuclei), which were also in the hold-out set of ecSeg-i, were used as a test set to prevent leakage. The remaining 297 images (22,970 nuclei) were used for ecSeg-c training/validation. For our NB patient tissue dataset, 260 of the 265 NB samples met our centromeric quality criterion, and were labeled by pathologists as ‘amplification’, or not. The NB patient tissue image dataset was split 75-25 into training/validation (5,350 nuclei from 193 images) and hold-out set (1,937 nuclei from 67 images).

### InterSeg architecture overview

Recall that interSeg has two distinct modules: ecSeg-i and ecSeg-c, both of which make predictions on individual nuclei annotated with DAPI and FISH. Therefore, we used an available method, NuSet^14^, to first segment each image into individual nuclei (Figure 2a). These individual nuclei are fed to ecSeg-i and ecSeg-c. Salient features and data filtering issues are described below, with details in methods.

**Figure 2:**
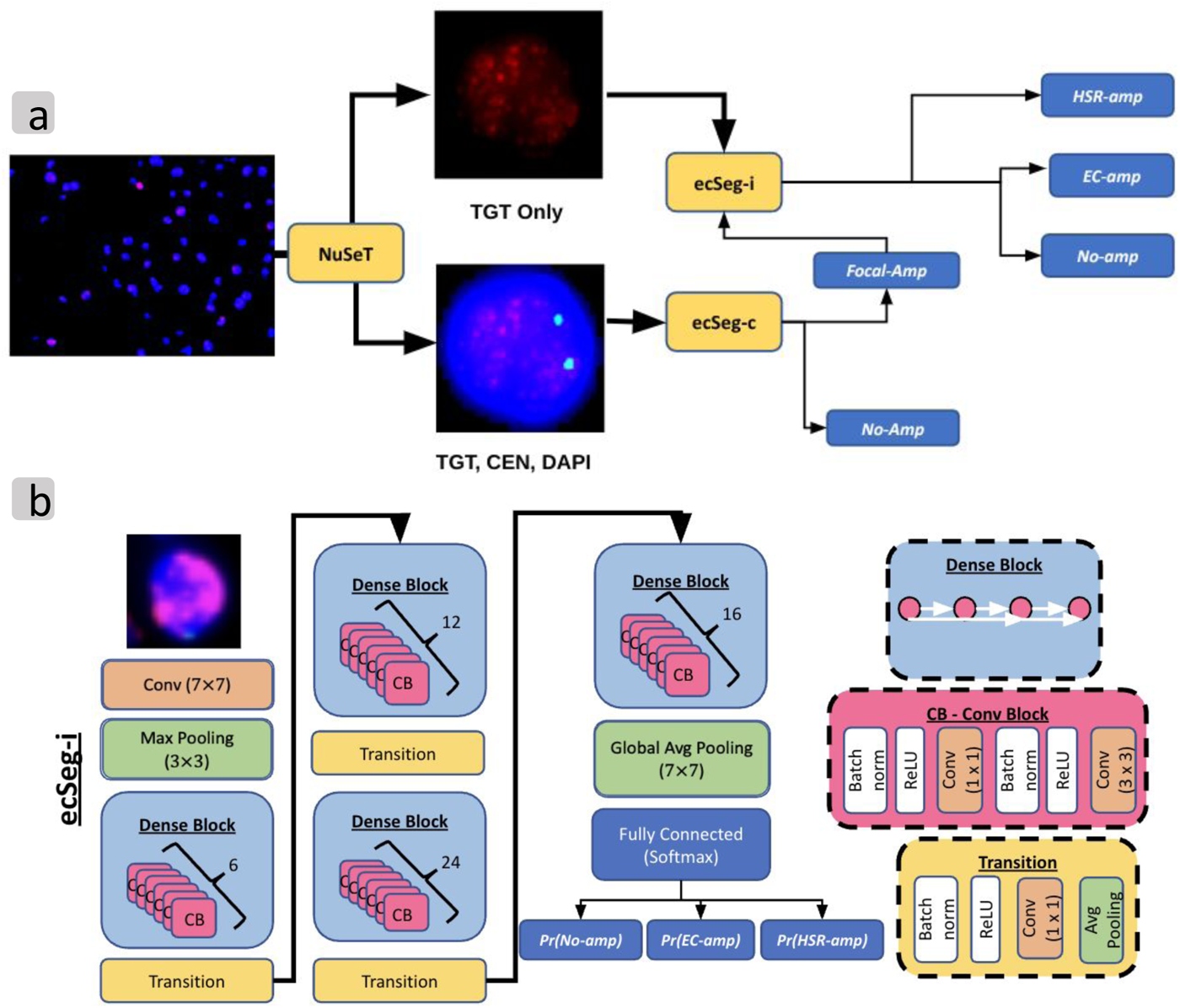
Tool pipeline. (a) InterSeg pipeline with 2 submodules: ecSeg-i and ecSeg-c. TGT: target FISH probe, and CEN: centromeric FISH probe. (b) EcSeg-i architecture based on DenseNet-121.

EcSeg-i and ecSeg-c are both based on the DenseNet-121 architecture^15^ (Methods). DenseNet-121 is a densely connected network with exhaustive skip connections between convolutional blocks, enabling feature-reuse throughout the network. The feature maps of all previous layers were concatenated and fed as input to the current layer, making it *densely* connected (Figure 2b).

The original DenseNet has a final classification layer with 1000 output nodes, corresponding to 1000 ImageNet^16^ classes. In our version of DenseNet for ecSeg-i, we used a final classification layer with 3 output nodes corresponding to the three output classes: EC-amp, HSR-amp, and no-amp. Notably, we also experimented with training a DenseNet from scratch and training with a network pre-trained on ImageNet. Although both networks achieved similar accuracy (F1-score difference of 0.1), the network trained from scratch converged much faster (∼50 epochs) than the pretrained network (∼80 epochs).

Ec-Seg-i returns the posterior probability of the membership in each class as the output for each nucleus. As a post-processing step, it optionally labels each nucleus with the amplification mechanism with the greatest posterior probability from ecSeg-i. Because there is considerable heterogeneity in ecDNA counts from cell to cell, most scientists make predictions based on groups of cells rather than individual cells. Therefore, we additionally generated cell line level metrics by employing a bootstrapping approach on the results obtained at the nucleus level (Methods). Briefly, this involved selecting 10 cells in each sampling instance and determining the most prevalent amplification mechanism within this group. This process was iterated 100 times, with a random selection of 10 cells in each iteration. The outcome of this iterative process was employed as the cell line level statistics.

EcSeg-i assumes that any amplification of the target is a focal event. However, amplifications can also occur due to aneuploidies, and this can be cytogenetically tested by using a centromeric probe. We trained a separate neural network (*ecSeg-c*) with the same DenseNet121 architecture as ecSeg-i (Supplementary Figure 2). For each nucleus, ecSeg-c predicts a binary classification label, focally amplified or not.

As with all deep-learning methods, a direct and intuitive explanation of the ecSeg-i posterior probabilities output is not available. This is specifically confounding for interphase FISH analysis where the high variability from cell to cell makes interpretation difficult even for the trained eye. To improve interpretation, we implemented a second module called *stat-FISH* to gather statistics that provide complementary evidence (Methods). These statistics are not used to change the output of ecSeg-i, but are provided as an addendum to ecSeg-i posterior probabilities. Importantly, in contrast to the per-cell posterior probabilities output by ecSeg-i, stat-FISH mimics human interpretation by analyzing and integrating the data from multiple nuclei (Supplementary Figure 3).

### Training and validation of interSeg modules

EcSeg-i converged on the training data after 50 epochs. We inspected what the architecture learned by visualizing the 7 × 7 filters of the first convolution layer and their corresponding feature maps over a test image. We observed that the majority of the filters initially learned to detect small circular objects, indicative of ecDNA patterns (Supplementary Figure 4). The corresponding feature maps show that the network is able to immediately separate the ecDNA-like structures from the background noise, affirming that the network is learning to recognize the object of interest.

We tested the robustness of interSeg predictions to variation of image acquisition, by artificially distorting the images (Methods), including shrinking, enlarging, and rotating (Supplementary Figures 5-7). In each case, the performance remained similar or identical to the non-distorted case. We also tested interSeg after changing contrast (Supplementary Figures 8-9), which can seriously impact intensity of the fluorescent signal. In a good image, we expect to see a bimodal distribution for the oncogenic FISH signal with a vast majority of pixels with very low intensity, and a small number of ‘true’ pixels with high intensity reflecting real probe hybridization. Lowering the contrast did not change the bimodality, but raising it led to significant bleeding of the FISH signal, impacting performance for HSR-amp lines but not EC-amp lines (Supplementary Figures 8-9). We used this result to generate a quality score for each image (Methods). Three of the tissue model test images were marked as low quality based on this method and were removed from final evaluation. Notably, the patient tissue samples were used only for hold-out testing of ecSeg-i. There were a total of 388 NB patient tissue images with pathologist annotations which were filtered for quality and for an annotation of ‘amplification’, or ‘no-amplification’ to yield 265 images used for testing of ecSeg-i.

We also observed some images with a weak but uniform centromeric signal (high kurtosis of mean nucleus centromeric intensity), in contrast with other images where there was a distinct centromeric signal, with high but varying intensity (low kurtosis). We filtered images with a kurtosis value greater than 3, and for those images ecSeg-c was not run, and only ecSeg-i was used to make calls. Additionally, we also defaulted to ecSeg-i when the maximum nucleus centromeric intensity was less than 10 (using a 0-255 scale), as these nuclei contain little centromeric signal. Of the 265 NB samples, 5 failed these centromeric quality score criteria, leaving 260 samples remaining. 67 were set aside as a hold-out test set for ecSeg-c, and 193 were used for training and validation.

### EcSeg-i and ecSeg-c accurately determine amplification mechanisms

In cases where a centromeric probe is not available, InterSeg defaults to running ecSeg-i (Figure 2a). Therefore, we tested ecSeg-i and ecSeg-c independently. EcSeg-i was tested on each of the 9,733 nuclei from the 118 cultured-cell and 97 tissue model images in the hold-out test data set. The 9,733 nuclei included 1,539 with no-amp, 3,497 nuclei with EC-amp, and 4,697 nuclei with HSR-amp. The model achieved F1-scores of 0.91 (recall: 0.91, precision: 0.91) for no-amp, 0.87 (recall: 0.91, precision: 0.84) for EC-amp, and 0.88 (recall: 0.86, precision: 0.91) for HSR-amp nuclei at the per-nucleus identification level (Figure 3a). These results are conservative estimates, as they assume uniform amplification within each cell line, despite the expected heterogeneity or lack of amplification in all cell lines.

**Figure 3:**
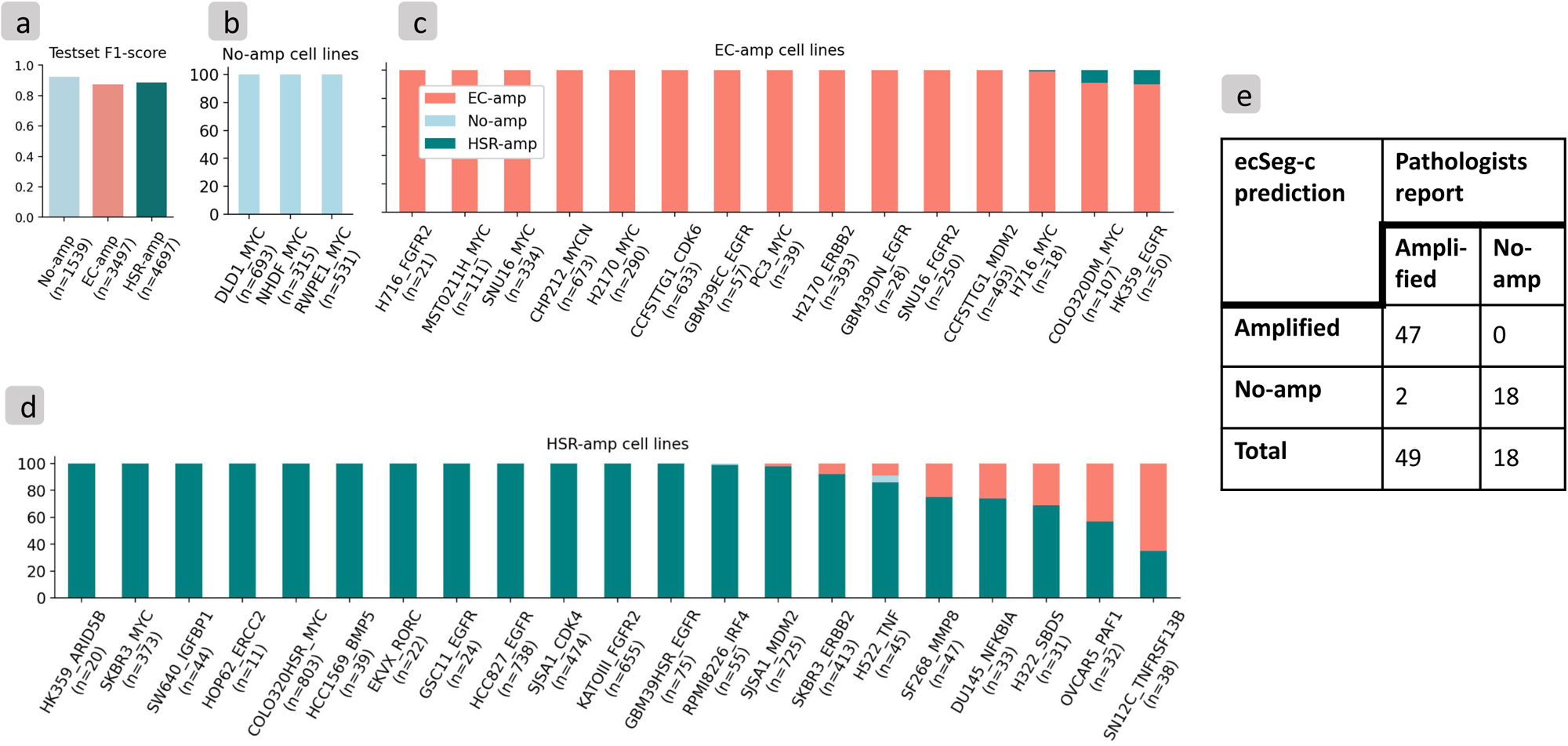
Testset results. (a) InterSeg F1-score on cultured cell line and tissue model test set, where n is the number of cells in each class. (b) Bootstrapped distribution of interSeg amplification mechanism of no-amp cell lines. (c) Bootstrapped distribution of interSeg amplification mechanism of EC-amp cell lines. (d) Bootstrapped distribution of interSeg amplification mechanism of HSR-amp cell lines. (e) EcSeg-c evaluation on NB hold-out set.

Next, we tested ecSeg-c. EcSeg-c was trained on the 297 tissue model images and 193 NB patient tissue images and subsequently tested on 95 tissue model images and 67 NB patient tissue images, as described earlier. On the tissue model images, we obtained a nucleus-level F1 score of 0.95 (recall: 0.95, precision: 0.94) and 0.99 (recall: 0.99, precision: 0.99) on the no-amp and amp classes respectively. For the NB patient tissue test subset, ecSeg-c obtained a 0.78 F1 score (recall: 0.92, precision: 0.67) on the no-amp class and a 0.92 F1 score (recall: 0.88, precision: 0.98) on the amp class at the per-nucleus level. For sample-level predictions (Figure 3e), we observed a 0.95 F1 Score (recall: 1.0, precision: 0.90) on no-amp and 0.98 F1 Score (recall: 0.96, precision: 1.0) on amp labels.

Notably, because we did not have metaphase annotations of ecDNA or HSR on the patient tissue images, the combined interSeg could only be tested on the cultured cells and tissue models. On the 118 cultured cell test images and 97 tissue model test images, interSeg obtained nucleus level F1 scores of 0.92 (recall: 0.97, precision: 0.88), 0.87 (recall: 0.91, precision: 0.85), and 0.89 (recall: 0.85, precision: 0.93) respectively for the no-amp, EC-amp, and HSR-amp classes.

The bootstrapped version of interSeg was tested 100 times on each of the 39 cell line-oncogene pairs to obtain single predictions for each pair. It correctly predicted the majority of the bootstrapped trials to carry the expected amplification in 38 out of the 39 samples overall (Figure 3b-d). Even in the non-bootstrapped nuclear level predictions, interSeg predicted more than 60% of the nuclei as EC-amp in all 15 ecDNA cell lines, more than 90% of the cells as no-amp in all 3 no-amp cell lines, and at least 50% of the cells as HSR-amp in 20 out of 21 HSR cell lines (Supplementary Figure 10 and Supplementary Table 2).

While bootstrapping eliminates small variability due to mis-prediction or noise, the remaining high variability in certain samples suggested heterogeneity between ecDNA and HSR. For example, the metaphase cell in SF268 shows two HSR amplifications. However, the stat-FISH data shows seven distinct FISH foci with a puncta pattern and an ecSeg-i posterior probability indicative of EC-amp (Supplementary Figure 11). In contrast, Supplementary Figure 12 shows a second SF268 nucleus with a high foci count of 9; in this case, ecSeg-i predicted the nucleus as primarily HSR-amplified due to the non-puncta distribution of the foci. Similar information can be found for SN12C (Supplementary Figure 13).

### InterSeg determines amplification heterogeneity between cell lines

To test interSeg prediction performance for heterogeneous samples containing both EC-amp and HSR-amp cells, we first created artificial composite images containing both ecDNA and HSR amplifications by combining the cells in the isogenic lines GBM39EC and GBM39HSR with the FISH probe *EGFR*. For the GBM39HSR cells in the computationally mixed images, we observed a 78%-22%-0% split between HSR-amp, ecDNA-amp, and no-amp predictions respectively. This mirrored the true GBM39HSR prediction percentages, which are 81%-19%-0% respectively. A statistical test could not distinguish between calls made on the pure HSR line versus the HSR labeled cells in the mixed image (chi-square test statistic: 2.3607, P-value: 0.1244). Similarly, for the true GBM39EC cells in the mixed images, we observed a 14%-86%-0% split between HSR-amp, EC-amp, and no-amp predictions respectively. Once again, this could not be statistically distinguished from the pure GBM39EC cell line, where the interSeg predictions were 14%-86%-0% HSR-amp, ecDNA-amp, and no-amp respectively (chi-square test statistic: 0.0186, P-value: 0.8914).

We repeated the experiment after concatenating pairs of test set images from COLO320DM and COLO320HSR with an absolute mean nuclei area difference of less than 50 pixels. This is a harder test because 29% of the cells in the used COLO320DM images were predicted to be HSR with a breakdown of 29%-71%-0% for HSR-amp, ecDNA-amp, and no-amp. Interestingly, the COLO320DM cells in the mixed images also showed a similar 32%-68%-0% distribution for HSR-amp, ecDNA-amp, and no-amp labels, respectively (chi-square test statistic: 1.7475, P-value: 0.1862). Similarly, observed predictions for COLO320HSR cells in the computationally mixed images were 97%-2%-1% for HSR-amp, ecDNA-amp, no-amp, respectively. These matched the interSeg predictions on pure COLO320HSR which were 96%-2%-2% respectively (chi-square test statistic: 0.3706, P-value: 0.8309).

Next, we also tested an experimental system where COLO320DM and COLO320HSR cells were grown on the same plate. An mCherry RFP tag was used to mark COLO320DM cells. A green DNA-FISH probe for *MYC* was used to test amplification in this mixed cell population (Methods). However, we also observed that the RFP tagging accuracy was not 100% and there were a small but unknown number of ecDNA cells that were not tagged (Supplementary Figure 14). Therefore, we utilized a latent parameter x denoting the number of COLO320DM cells that were not RFP tagged. Next, we computed the optimal value for x that maximized the likelihood of the observed frequencies seen in the pure cell line test datasets for COLO320DM and COLO320HSR (Methods). At x=6.14% (which would imply a tagging accuracy of 93.86%), we observed a strong correlation, or no statistically significant difference between expected heterogeneity and observed heterogeneity (chi-squared test statistic: 2.7252, P-value: 0.4360). Together, these results illustrate the power of interSeg in predicting amplification mechanisms in the presence of heterogeneity.

Based on these results, we decided to use the following rule based on predictions after bootstrapping: A cell line was considered to be no-amp at least 80% of the cells were classified as no-amp; as HSR-amp if at least 80% of the cells were classified as HSR-amp; as EC-amp, if at least 50% of cells were EC-amp. Otherwise, the sample was classified as mixed or heterogeneous.

### stat-FISH provides an explanation of amplification status

Because interSeg uses deep neural networks to determine the amplification mode, there is limited insight into the features used to make this decision (for partial information, see Supplementary Figure 4). Therefore, we analyzed the data with stat-FISH, a complementary module that computes statistics of the distribution of oncogenic foci per cell. As expected, cells with EC-amp had higher mean and variance in the copy number signal compared to HSR-amp cells. Only 40% of the EC-amp images had a mean < 10 and variance < 64, in contrast to 97% of HSR-amp images with those properties (Figure 4a and Supplementary Table 4). The number of foci and the total FISH signal were also significantly higher in EC-amp cells, whether analyzed across all cell lines (Figure 4b,c) or for individual pairs (e.g., Figure 4d and Supplementary Tables 5, 6, 7). Despite these differences, there was high variability in the number and spread of FISH foci across samples. The maximum accuracy of stat-FISH amplification status prediction on the test samples, across different cut-offs of mean and variance, was 83%, lower than the 97% sample accuracy of interSeg (Supplementary Table 8 and Methods). Thus, while stat-FISH is a useful complementary method that allows for an intuitive understanding of amplification modes, it lacks the predictive power of the deep neural network of interSeg, which may be correcting for signal-to-noise ratio, changing morphologies, and latent correlations.

**Figure 4:**
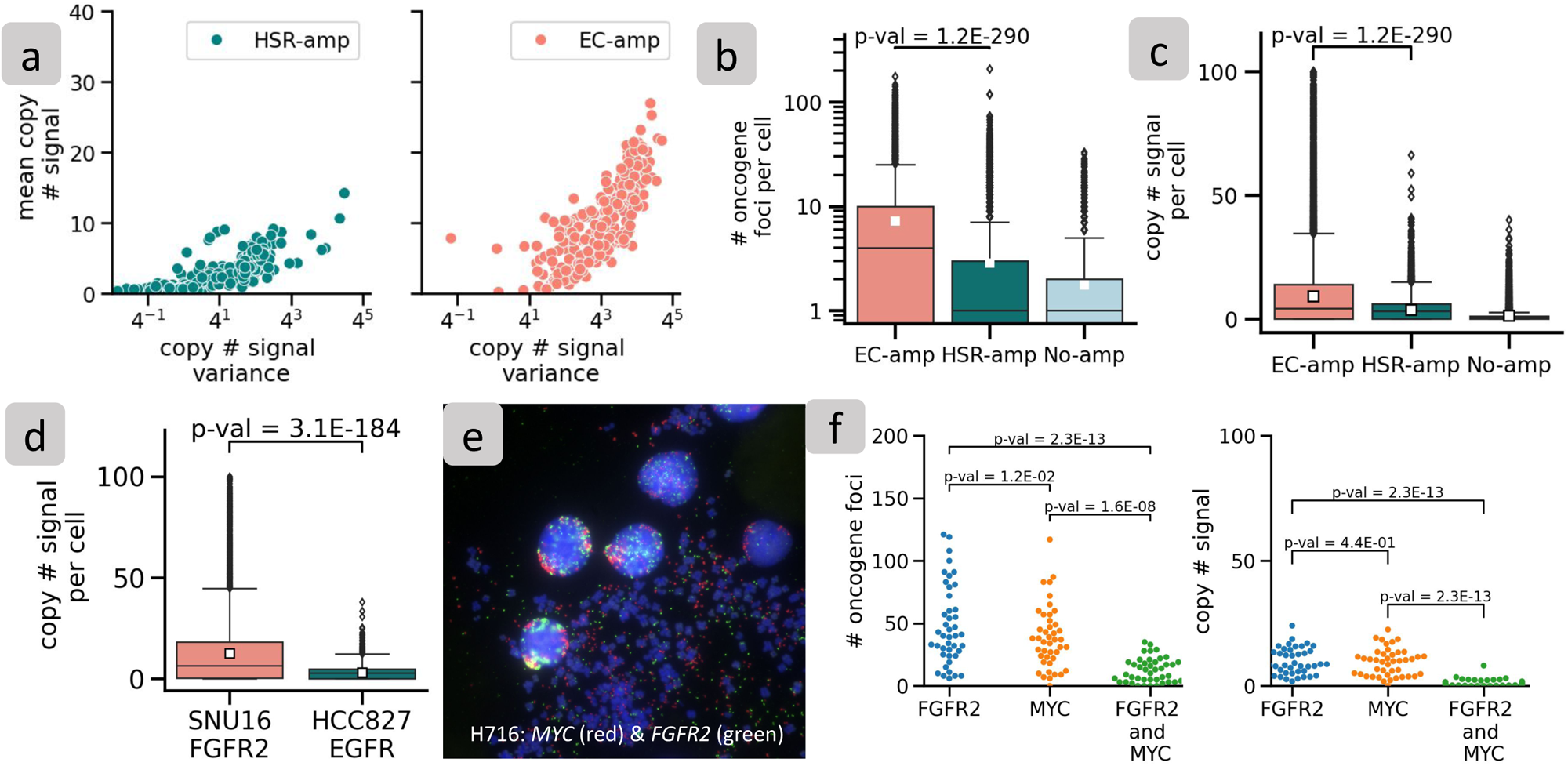
Explaining amplification mechanism using Stat-FISH. (a) Image-level mean and variance of copy number signal for HSR-amp and EC-amp images. (b) Number of oncogene foci per cell across all cell lines separated by EC-amp, HSR-amp, and no-amp. (c) Copy number signal per cell across all cell lines separated by EC-amp, HSR-amp, and no-amp. (d) Copy number signal of an EC-amp cell line (SNU16) and a HSR-amp cell line (HCC827) (e) Example of probing multiple oncogenes within a metaphase/interphase spread in H716 (*MYC* red, *FGFR2* green). (f) Number of oncogene foci and copy number signal of *FGFR2* and *MYC* oncogene for each cell in H716.

### stat-FISH allows for exploratory quantification of multiple oncogenes

While stat-FISH cannot predict amplification status with as much accuracy as interSeg, it nevertheless provides the flexibility for additional computations that are not easy with interSeg. For example, we used stat-FISH to investigate H716, a colorectal cancer cell line where interSeg predicted EC-amp for two distinct probes, corresponding to *FGFR2* and *MYC*, for each investigated cell (Figure 4e and Supplementary Table 3). The included metaphase in the figure confirms the correctness of the two predictions as being distinct ecDNA. We next quantified the FISH signal using stat-FISH. The average and median copy number signal for *FGFR2* was at 10 and 9, respectively, similar to those of *MYC*, which were 9 and 10, respectively. Similar to metaphase, we also found examples of co-occurring *FGFR2* and *MYC* amplification signals that showed up as yellow (*FGFR2*-green and *MYC*-red). The stat-FISH results suggested a higher count of *FGFR2* ecDNA relative to *MYC*, and the individual numbers were significantly higher than co-occurrences (Figure 4f). However, the co-amplification signal was also strong, and significantly higher relative to chance occurrence (Mann-Whitney U test P-value 1.6E-08; Methods). This result suggests either that the ecDNA species interact^17,18^ or the existence of ecDNA that carry both *MYC* and *FGFR2*.

### InterSeg determines heterogeneity of ecDNA in patient tissue samples

Across the 265 patient tissue NB samples, the interSeg predictions were 167 EC-amp (63%), 65 as no-amp (25%), and 33 heterogeneous (12%). These samples were previously classified by pathologists as ‘amplification’ or not (Figure 5a), where ‘amplification’ included the EC-amp, HSR-amp, and heterogeneous calls made by InterSeg. Among the 71 pathologist annotated ‘no amplification’ samples, interSeg labeled 63 (89%) as no-amp, 5 (7%) as heterogeneous, and 3 (4%) as EC-amp. When limited to the test samples, interSeg called 15 of 18 (83%) as no-amp, and 3 (17%) as heterogeneous (Supplementary Figure 16). Similarly, among the 194 pathologist annotated ‘amplification’ category, interSeg labeled 164 samples (85%) as EC-amp, 28 (14%) as heterogeneous, only 2 (1%) as no-amp. When limited to the test samples, interSeg called 41 of 49 (84%) as EC-Amp, 7 as heterogeneous (14%), and only 1 (2%) as no-amp.

**Figure 5:**
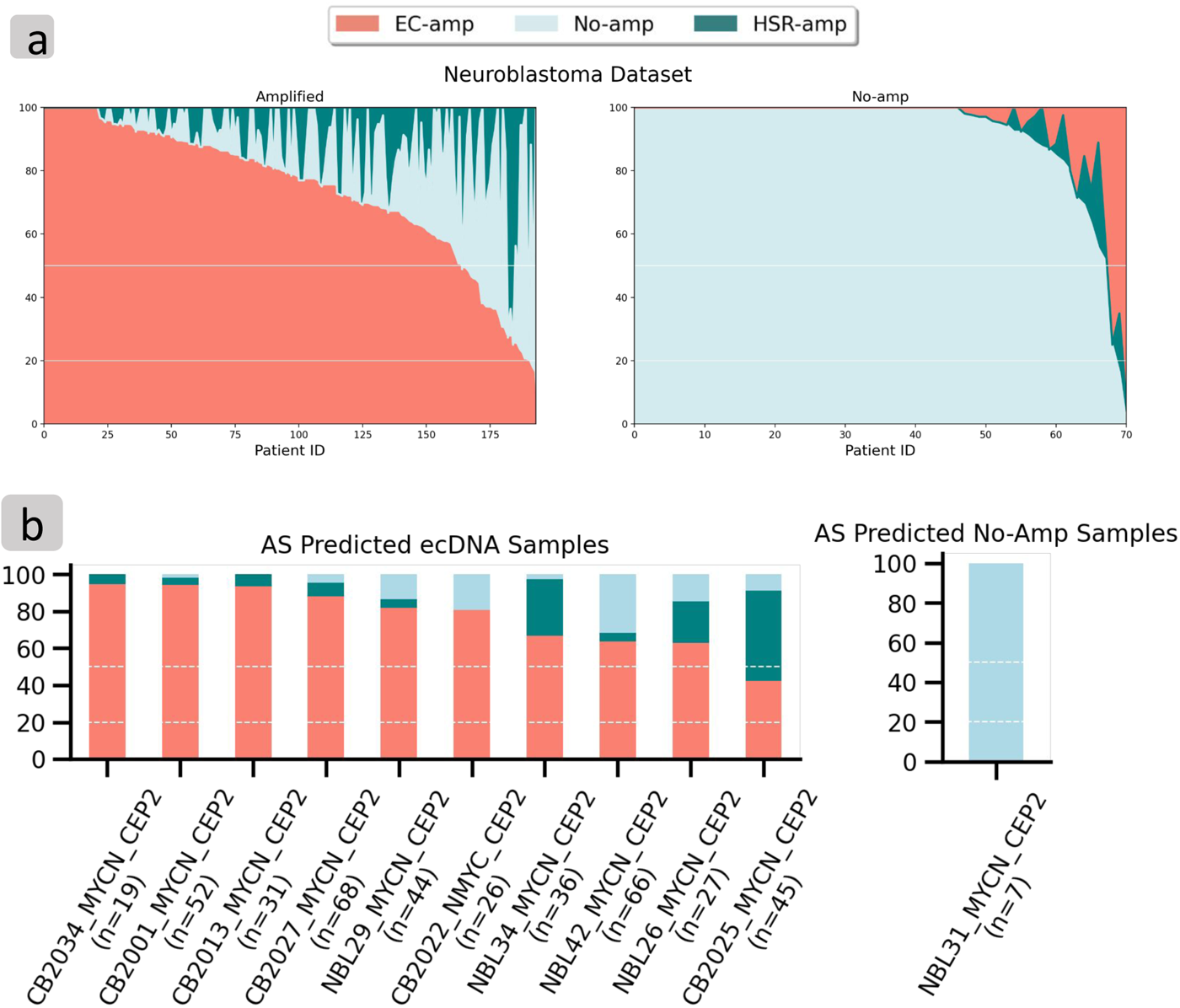
Patient Tissue Results. (a) Non-bootstrapped distribution of interSeg amplification mechanisms across all NB samples with pathologist annotation. The samples were stratified by ‘no-amplification’ and ‘amplification’ labels annotated by pathologists. Each column corresponds to a single patient, and the bar height corresponds to the proportion of cells labeled for each amplification class by interSeg. (b) InterSeg predictions on NB test set samples compared with calls based on Amplicon Suite analysis of Whole Genome Sequencing Data.

Because the pathologists used a binary classification between focally amplified or not, we also tested the majority call. On the hold-out NB test data from no-amplification category, interSeg called the majority class as no-amp in 18 (100%) of 18 samples. Similarly, in the focal amplification test data, interSeg called the majority class as amplified in 46 (94%) of 49 samples. Moreover, 43 of the 46 focal amplification calls were labeled as EC-amp, consistent with the high prevalence of ecDNA in *MYCN* amplified neuroblastoma samples^19^. Thus, the interSeg results were highly consistent with pathologist annotations, but provided additional information in terms of cellular heterogeneity, and mode of amplification.

To further validate interSeg’s prediction accuracy, we gathered a set of hold-out NB test samples where whole genome sequencing (WGS) data had been acquired. After filtering for quality score and removing samples with fewer than 5 nuclei, 11 NB samples remained (Supplementary Table 11). The Amplicon Suite pipeline (AS) is routinely used to identify ecDNA using WGS^6,20^. Upon analyzing the WGS data with AS, 10 samples were predicted to be cyclic, indicative of ecDNA containing *MYCN*, and 1 showed no focal amplification.

9 of the 10 AS predicted ecDNA samples were also predicted by interSeg to be EC-amp (Figure 5b). For the one remaining AS predicted ecDNA sample, interSeg predicted heterogeneity, with 49% of nuclei as HSR-amp, 42% of nuclei as EC-amp, and 9% of the nuclei as No-amp. The results are highly concordant, because AS makes a single call based on bulk sequencing, and ecDNA have been previously observed to reintegrate into a non-native chromosomal location. One sample was predicted by AS as carrying no focal amplification, and interSeg predicted 100% of the nuclei as no-amp.

## DISCUSSION AND CONCLUSION

Cytogenetically identifying the amplification mechanism in interphase cells is an important and incompletely understood problem. Although sequence-based methods can reconstruct focal amplifications, they cannot fully capture the dynamic nature of ecDNA and the amplification mechanism of a cell’s *present* state. Image-based tools can accurately reconstruct ecDNA in fluorescently stained images of cells in metaphase in which the ecDNA is clearly visible as tiny DNA particles floating separately from the chromosomes. However, this requires sampling of cultured or mitotic cells, and is difficult to obtain from patient tissue images. Patient tissue images primarily contain densely clustered interphase cells, where the DNA is inside an intact nuclear membrane and loosely wound. Moreover, ecDNA counts vary from cell to cell and include many cells with low counts. This makes it extremely challenging to discern ecDNA even for a trained eye. Nevertheless, on multiple data sets including cultured cells, tissue models, and patient tissue, and on experimentally and computationally mixed cells containing both ecDNA and HSR amplification, interSeg was able to predict heterogeneity accurately, and works well in models of tissue slices.

We also present a companion method, stat-FISH, that provides interpretability to interSeg results and provides useful statistics for deeper analysis. We demonstrated various use cases of interSeg+stat-FISH in predicting amplification status, amplification heterogeneity between EC-amp, HSR-amp, and no-amp cell lines, and reconstructing the amplification profile of multiple oncogenes within a single cell. Most importantly, we show that interSeg accurately quantifies the amplification mechanism of patient tissue images.

InterSeg is flexible enough to use without a centromeric probe, but we recommend using it with a centromeric probe. Also, it is run in an optional bootstrap mode, which smooths the nuclei results through a voting mechanism. The bootstrap mode is best utilized in situations where no heterogeneity is expected, and a single label can be applied to the entire image. In the presence of heterogeneity, interSeg should be run in a non-bootstrapped mode. In the manuscript, we utilized the non-bootstrapped mode for analysis of patient tissues and the heterogeneity experiments, but provided data on both bootstrapped and non-bootstrapped runs.

Even though we use a unified model for cultured cells, tissue models, and patient tissue, the three modes are quite different. Especially, patient tissues often contain multiple cell types, including normal cells. Currently, interSeg does not correct for tumor purity. Additionally, it uses a third party tool to separate individual nuclei, but may not be able to adequately separate tightly packed nuclei, which in turn could influence the predictions of the number of FISH foci, and the amplification mechanism. In future work, we will experiment with interphase cultured cells where cells are cultured on a cover slip and are not perturbed by any chemicals, to better match the nuclear distribution of ecDNA on patient tissues. InterSeg corrects for problems due to contrast and other quality control issues but more data will be needed to understand the degradation of performance on lower quality data sets. Another challenge is that most pathologists do not distinguish between HSR and EC-amp making it difficult to experimentally validate the amplification mode (ecDNA/HSR) on patient tissue. Moreover, whole genome sequencing signatures from bulk cells also cannot distinguish ecDNA from HSRs formed by reintegration of ecDNA into the chromosomes while maintaining their structural features. These are all areas of development which will be addressed in ongoing and future work. Despite these challenges, interSeg has been successfully applied to hundreds of patient tissue images, and should be a useful tool for analysis of focal amplification mechanisms.

## METHODS

### Image Acquisition Protocols

To generate the cultured cell images, we arrested the cells by treating them with Colcemid (Karyomax) at a final concentration of 0.1 μg/mL for 1-5 hours. Cells were collected, washed with PBS, and re-suspended in 75 mM KCl for 10-15 minutes at 37 °C. The hypotonic buffer reaction was quenched by adding an equal volume of Carnoy’s Fixative (3:1 Methanol:Glacial Acetic Acid). Cells were centrifuged, washed, and re-suspended in Carnoy’s fixative three more times. They were then re-suspended in 100-400 μL of Carnoy’s Fixative and dropped onto non-overlapping sections of humidified slides. Slides were equilibrated in 2x SSC and dehydrated in an ascending alcohol series of 70%, 85%, and 100% ethanol for two minutes each. The appropriate DNA FISH (Empire Genomics) probe was added to the sample and placed on a 75 °C slide moat for 3-5 minutes to melt the DNA. Probe hybridization occurred at 37 °C in a humidified slide moat for 4 hours to overnight. Slides were washed for two minutes each in 0.4x SSC and 2x SSC/0.1% Tween-20. Slides were stained with DAPI, washed in 2x SSC and ddH2O, and then mounted with mounting media (ProLong Gold or Vectashield). Cover slips were sealed with clear nail polish to prevent drying of the sample. Images were captured using a 63x objective on either an Olympus BX43 wide-field fluorescent microscope or a Leica Thunder Imager.

The tissue model images were derived from xenografts. The CytoCell Tissue Pretreatment Kit (LPS 100, Oxford Gene Technology IP Ltd.) was used for heat pretreatment of Formalin-Fixed, Paraffin-Embedded (FFPE) tissue prior to Fluorescence In Situ Hybridization (FISH). All FISH probes were purchased from Empire Genomics Inc. FFPE slides were baked at 50 °C overnight, deparaffinized three times with xylene (1330-20-7, Millipore Sigma) for 10 minutes each, and immersed in 100% and 70% ethanol (64-17-5, VWR International LLC) for 2 minutes each. After washing in water for 2 minutes, the slides were incubated in a pretreatment solution at 100 °C for 40 minutes. Slides were dehydrated in a graded ethanol series of 70%, 85%, and 100%, and then air-dried. Next, 10 μL of probe mixture was applied to the hybridization area, cover-slipped, and sealed with CytoBond coverslip sealant (2020-00-1, SciGene Corp.). Slides were incubated in the ThermoBrite System (Abbott) at 80 °C for denaturation and hybridized at 37 °C for 16 hours. After gently removing the coverslip sealant, the slides were immersed in 2x SSC/0.1% Tween-20 (V4261, Promega Corp.) for 3 minutes in the dark. The coverslips were slipped off the slides while still in the SSC buffer. Next, slides were washed in 0.4x SSC solution at 73 °C for 2 minutes, transferred to water for 1 minute, air-dried in darkness, and stained with DAPI (DFS500L, Oxford Gene Technology IP Ltd.), and cover-slipped. FISH results were examined with a Keyence fluorescence microscope (BZ-X800 model, Keyence Corp.).

### Data pre-processing

We pre-processed the images by delineating each individual intact nucleus in the image. We used a package called NuSeT^14^ to identify and segment each nucleus. NuSeT utilizes multiple neural networks to identify and separate each nucleus, even in dense, overlapping clusters. We drew a bounding box around each unique nucleus, cropped the region to the bounding box, and resized the crop to a 256 by 256 patch (Figure 2a). For bounding boxes larger than 256 by 256 pixels, we applied a sliding window approach to obtain multiple 256 by 256 patches, with each patch analyzed seperately.

For the ecSeg-c model, each input patch contains channels corresponding to the DAPI probe, centromeric probe, and target probe. To control for the variation in brightness between channels, we uniformly rescaled the DAPI channel to the range 0 to 1. Additionally, we jointly rescaled the target and centromeric probe to the range 0 to 1.

### EcSeg-i and ecSec-c architecture

The backbones of ecSeg-i and ecSeg-c are the DenseNet-121 architecture^15^. Densenet-121 is a 121 layered convolutional neural network (CNN) with 12 layers. The feature maps of all previous layers are concatenated and fed as input to the current layer, making it *densely* connected. The primary benefit of this dense connection is that it enables deeper layers to reuse features learned in earlier layers without having to relearn them. Consequently, a DenseNet uses fewer parameters than an equivalent vanilla CNN.

EcSeg-i and ecSeg-c are composed of four dense blocks containing, 6, 12, 24, and 16 convolutional blocks, respectively. Each convolutional block is composed of 6 sequential operations: batch normalization (BN), a rectified linear unit (ReLU), 1 × 1 convolution, BN, ReLU, and a convolution. The dimensions of all the feature maps within a dense block are kept the same (i.e. no down-sampling) but the number of filters increases by a growth factor 𝑘 = 32. This makes it practical to concatenate the feature maps instead of summing them.

Each convolutional block adds 32 additional feature maps. In total, DenseNet-121 has one 7 × 7 convolutional layer, 58 3 × 3 convolutional layers, 61 1 × 1 convolutional layers, 4 averaging pooling layers, 1 max pooling layer, and one fully connected layer.

The original DenseNet-121 used a final classification layer containing 1000 output nodes as it was trying to classify 1000 classes. For the ecSeg-i model, we use a final classification layer containing 3 output nodes corresponding to the three output classes: EC-amp, HSR-amp, and no-amp. For the ecSeg-c model, we use a final classification layer with 2 output nodes, corresponding to the no-amp and Amp output classes.

### Training Procedure

We trained both the ecSeg-i and ecSeg-c models on 4 GeForce GTX 1080 Ti GPUs using the Adam optimizer. For ecSeg-i, we used a patience criterion of 7, and a learning rate of 1e-4. If the validation loss did not improve for 7 epochs the training was halted. We minimize the cross-entropy loss function to train our network. We trained the network for 200 epochs and found that the model converged after 120 epochs. To find the optimal architecture, we performed grid-search of the following hyperparameters.

When training the ecSeg-c model, we initialized the network with the DenseNet121 weights pretrained on the ImageNet dataset. We used a patience criterion of 7 epochs, with the validation set area under the curve (AuC) metric as the early stopping criterion. Additionally, we used the Adam optimizer with a learning rate of 5e-4, and minimized the binary cross-entropy loss. To control for class imbalance, we applied balanced sampling during each epoch across each tissue type (tissue model/patient tissue) and amplification type (no-amp/amp) pair. We assigned equal sampling weight to each of the 4 tissue type pairs, effectively downsampling the majority classes. We trained the ecSeg-c network for 200 epochs and observed the model converge after 18 epoch. We noted that less training epochs were needed for convergence due to the use of pretrained initial weights. Similar to ecSeg-i, we performed a grid-search for the used hyperparameters.

### interSeg bootstrapping of cell predictions

We note that each experiment typically contains several images. For example, Figure 3b contains 933 viable cells across 6 images of the DLD1 cell line with *MYC* staining. Similarly, we found 368 viable cells across 6 images of the NHDF cell line with *MYC* staining. To address variation in cell and image counts, we applied a custom bootstrapping approach as follows. First, we assigned a single amplification mechanism for each cell based on the highest likelihood prediction from interSeg. Next, our procedure randomly selected 10 cells and identified the predominant amplification mechanism based on the majority vote. We iterated this process 100 times to produce a distribution representing the overall amplification mechanism of the experiment. The original interSeg calls are also retained for comparison, etc.

### Quality Score Filtering

For each image, we generate an oncogenic probe quality score which indicates whether the image is apt for interSeg. We first bin the oncogenic FISH signal into 50 buckets based on their pixel intensities. We then find the highest peak left of the 25th bin and right of the 25th bin. We find the peaks by simply comparing the neighboring values. We compute the quality score, 𝑄, by dividing the leftmost peak (ℎ_1_) by the rightmost peak (ℎ_2_), 𝑄 = ℎ_1_/ℎ_2_. Images with 𝑄 < 0. 2 were marked as low quality. Additionally, we excluded nuclei with a mean oncogenic FISH signal below 0.05 from both the interSeg and ecSeg-c analyses, as these nuclei exhibited extremely low oncogenic FISH signal.

We generated a centromeric probe quality score for each image as well, based on the kurtosis of the mean centromeric intensity per nucleus. Images with a kurtosis value greater than 3 were marked as having low centromeric probe quality and were excluded from ecSeg-c analysis, defaulting to evaluation in interSeg target-channel-only mode. Additionally nuclei with maximum centromeric pixel intensity less than 10 were also excluded from ecSeg-c analysis and defaulted to the ecSeg-i prediction.

### stat-FISH Segmentation Preprocessing

We employed instance segmentation to decrease the occurrence of overlapping nuclei receiving an inflated foci count. We utilized the min-cut algorithm to transform the binary segmentation output from NuSeT into an instance segmentation. To separate overlapping nuclei, we generated a 4-connectivity pixel graph for each connected component in the NuSeT segmentation. To identify nucleus centers, we applied an L1 distance transformation to the NuSeT segmentation and selected local maximums with greater than a 10-pixel distance away from the nearest background pixel. For a given connected component, we determined the minimum number of edges to be removed to isolate two centers in the pixel graph. We applied this algorithm to all connected components exceeding 1.25 times the median connected component area and separated them based on a flow limit of 60. This preprocessing method was only used for stat-FISH and not interSeg, since stat-FISH outputs a quantitative rather than categorical prediction.

### stat-FISH

Stat-FISH looks for local peaks in brightness in the FISH channel. The input for stat-FISH is a single 8-bit FISH-probe channel, and its corresponding binary image representing the nuclei segmentation from NuSeT. To establish whether a given pixel is classified as an foci (local peak), we have three criteria:

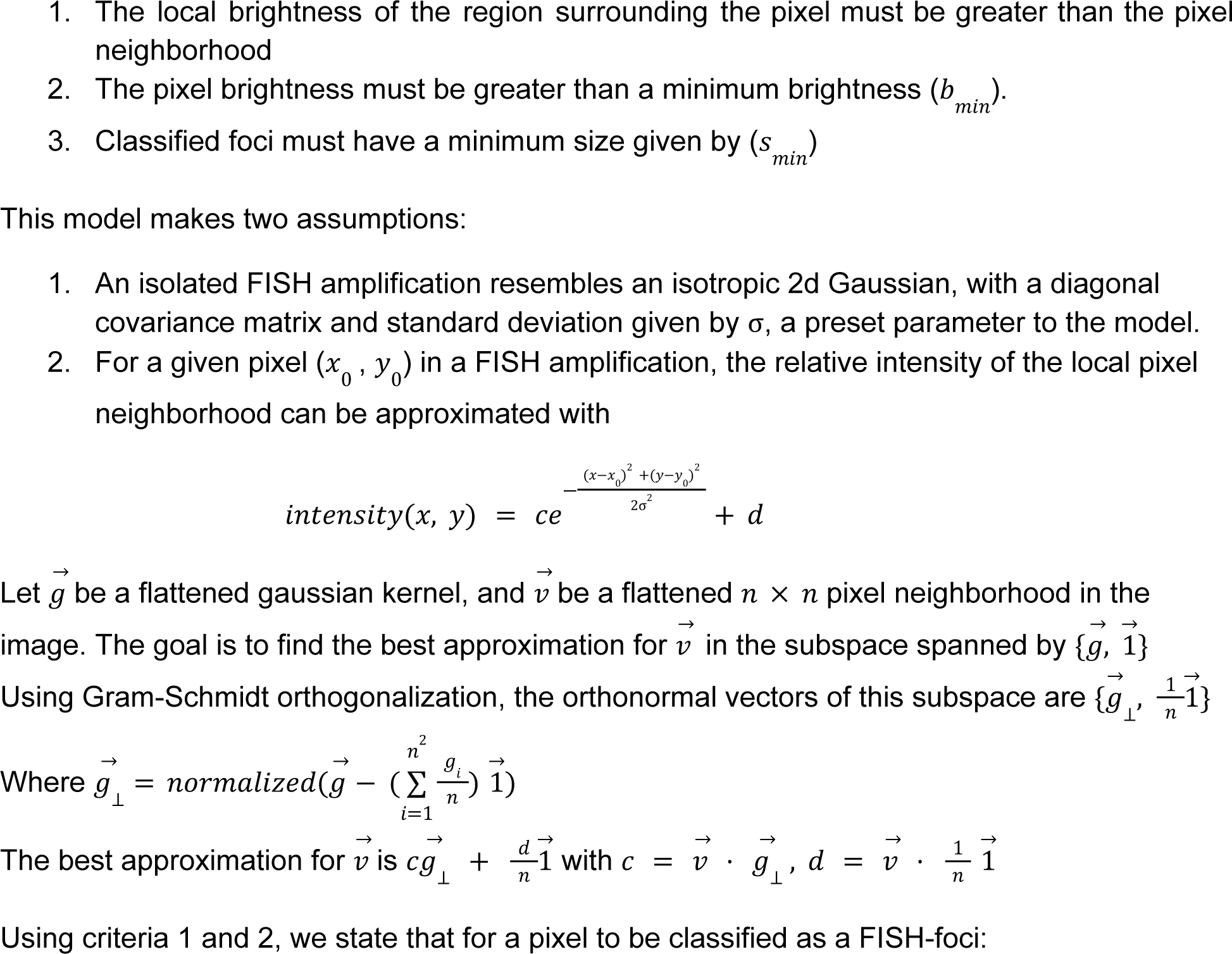
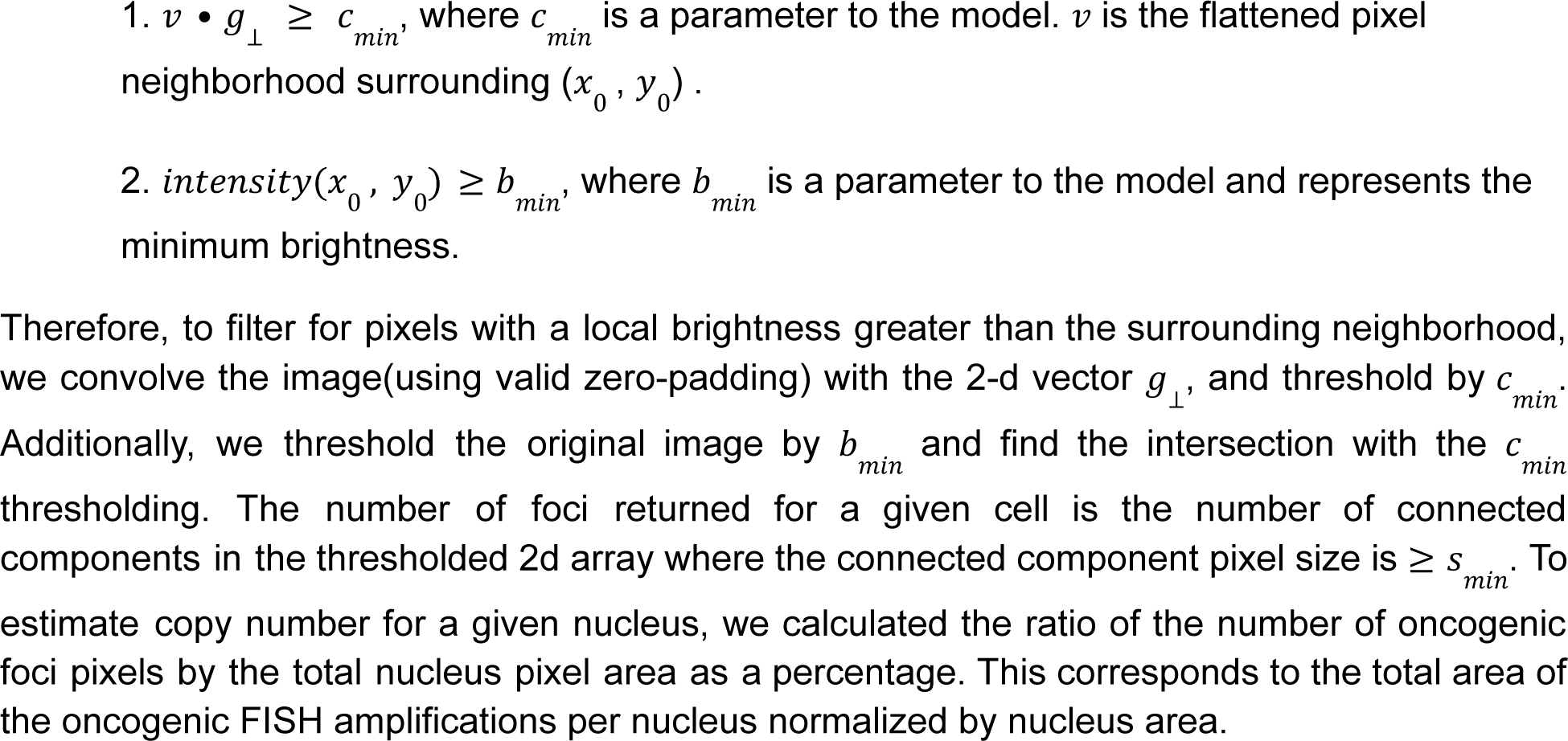

We emphasize that stat-FISH is a deterministic tool and primarily quantifies the visual data in the image. It does not accurately determine the amplification mechanism in every case. However, it is useful to understand the interSeg predictions, and is used in conjunction with interSeg.

### Image distortion

We tested the robustness of interSeg by testing against distorted images, including enlarging, shrinking, rotating, and modulating their contrast (Supplementary Figures 5-7). We chose one image from an EC-amp cell line and one from an HSR-amp cell line. For each image, we shrunk them by 80%, enlarged to 1.2x the original size, rotated them 45 degrees, decreasing the contrast by 40%, and increasing the contrast by 40%.

### Red Fluorescence Probe tagging for generating heterogeneous samples containing ecDNA and HSR

The COLO320DM and COLO320HSR cell lines used in the study are clones with comparable *MYC* copy numbers, selected from cells obtained from ATCC. COLO320DM H2B-mCherry was engineered by lentiviral infection of H2B-mCherry into isogenic COLO320DM cells, followed by sorting of mCherry-positive cells. Two rounds of cell sorting were performed to ensure that about 95% of the COLO320DM H2B-mCherry line were mCherry-positive.

Both COLO320HSR and COLO320DM H2B-mCherry were cultured in DMEM supplemented with 10% FBS and penicillin-streptomycin. One million cells from each cell line were harvested at 70%-80% confluency and fixed with 4% paraformaldehyde for 10 minutes at room temperature, followed by two washes with 1x PBS. The fixed COLO320HSR and COLO320DM H2B-mCherry cells were mixed at a 1:1 ratio and cytospun onto an imaging slide at 800 rpm for 8 minutes using the low mode on a Thermo Scientific Cytospin 4 Centrifuge.

Immunofluorescence (IF) of mCherry was performed on the slide to distinguish between COLO320HSR and COLO320DM H2B-mCherry cells. DNA FISH targeting *MYC* (https://empiregenomics.com/product/16399) was performed following IF to detect *MYC* amplification. Images were acquired with a Leica SP8 LIGHTNING confocal microscope.

### Estimation of RFP Tagging Accuracy in Hybrid COLO320 Experiment

Our goal in this analysis is to validate that the observed frequencies of predicted annotations and mCherry status (tagged vs not tagged) match our expectations. The categories for each nucleus are:

a. mCherry tagged and EC-amp predicted
b. not mCherry tagged and EC-amp predicted
c. mCherry tagged and HSR-amp predicted
d. not mCherry tagged and HSR-amp predicted

To determine whether a nucleus is mCherry tagged, we took the maximum mCherry brightness over all of the pixels in the segmented nuclei. We classify a nucleus as mCherry tagged if its maximum pixel intensity exceeds 10 pixel brightness. We selected the threshold of 10 pixel brightness as it strikes a balance between precision and recall in the AuC Curve for both

HSR-amp and EC-amp cells (Supplementary Figure 15, Supplementary Table 14). Since we used maximum pixel intensity as the threshold, we excluded nuclei near the boundary, as pixels from these nuclei are missing.

Since mCherry is an RFP which is inserted into COLO320DM, we expect only the ecDNA-amp nuclei to be tagged, given 100% mCherry tagging accuracy.

However, the mCherry tagging accuracy is likely lower than 100%, and this analysis aims to estimate the number 𝑥 of true ecDNA nuclei that are not mCherry tagged.

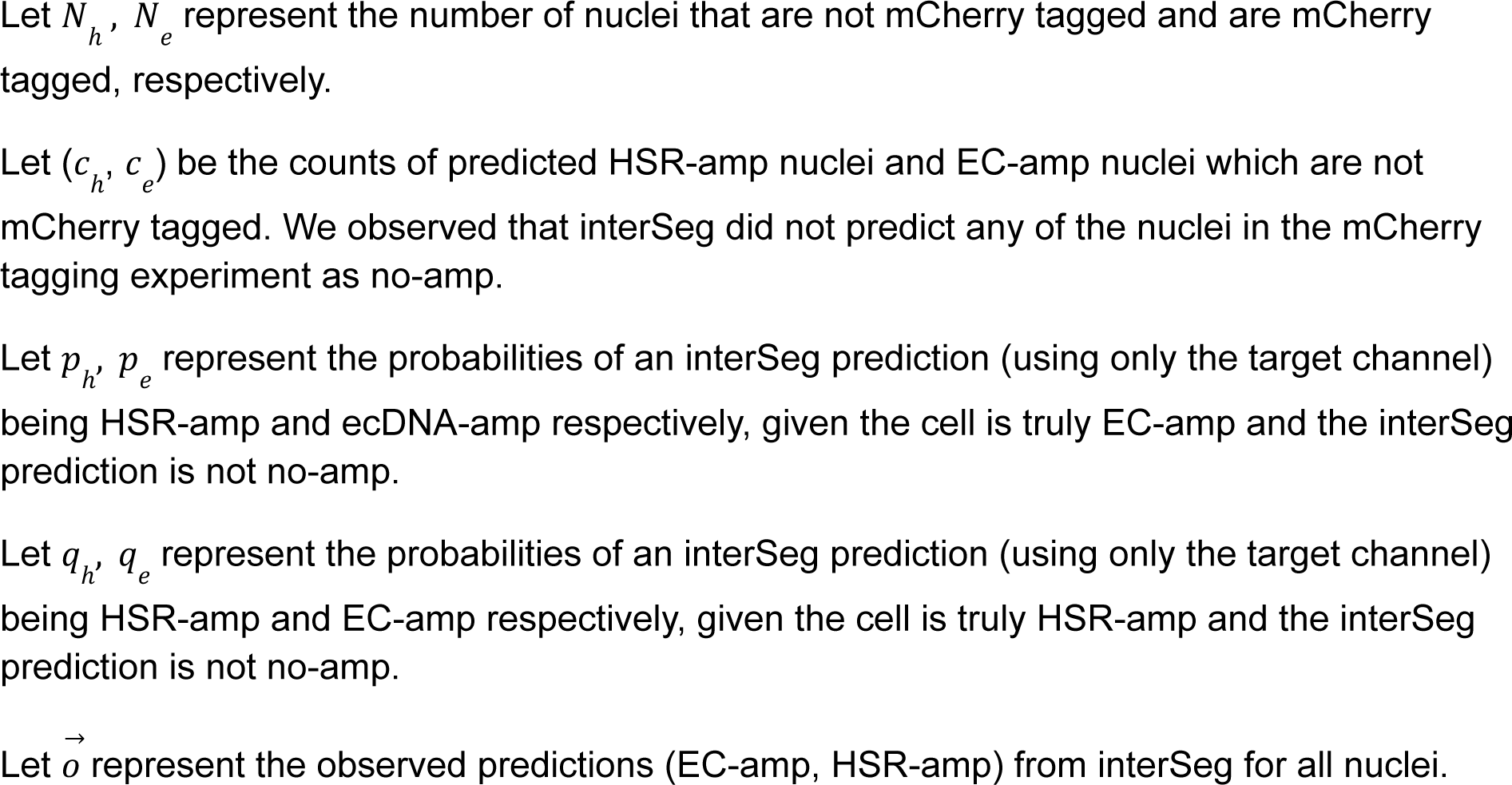
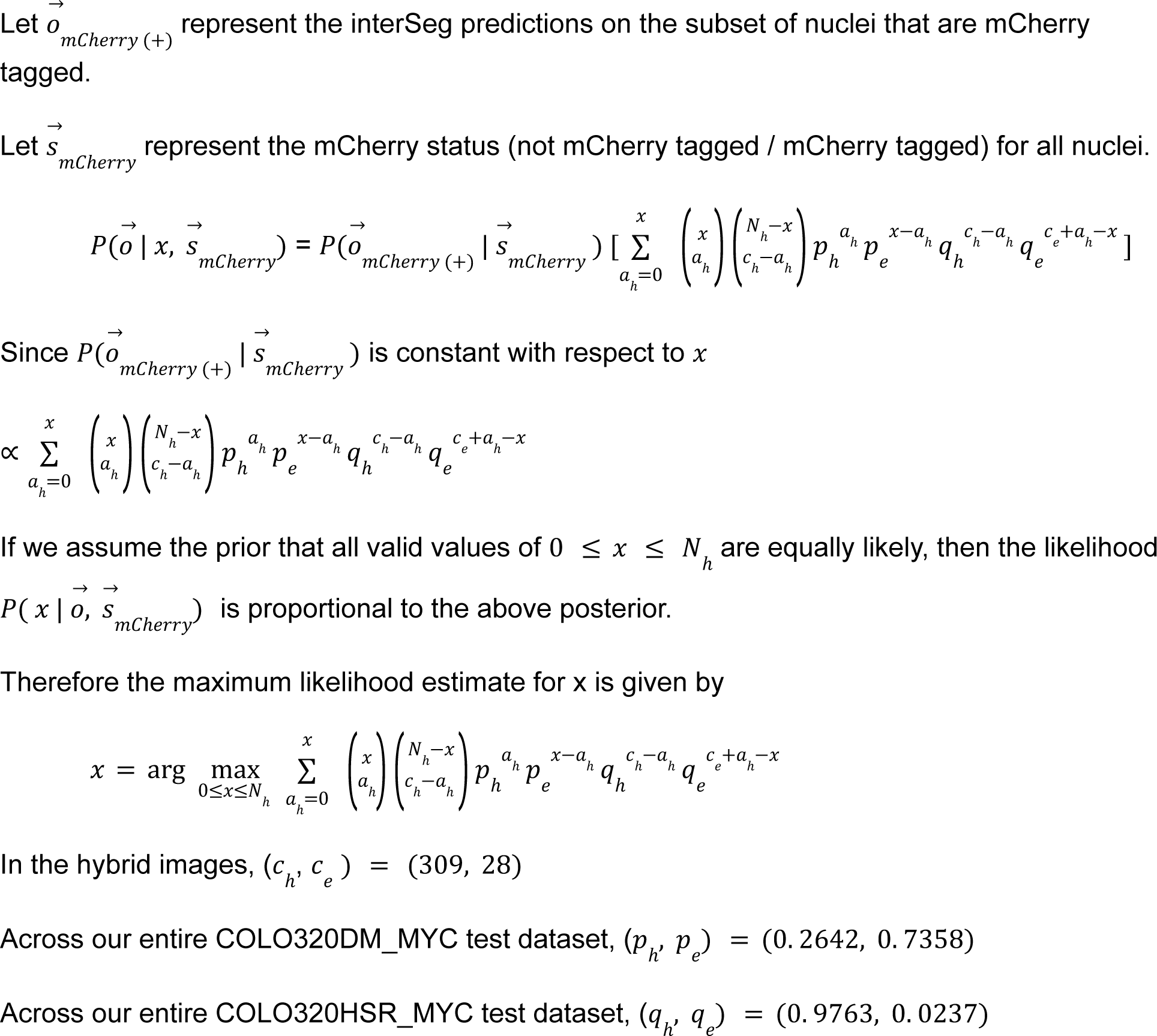

The value x, which represents the number of EC-amp nuclei which are not mCherry tagged, maximizes the above likelihood when x=28 nuclei.

To test this x=28 nuclei prediction, we performed a chi-squared test on the expected vs observed nuclei counts on the 4 categories (mCherry tagging status and interSeg prediction). With a chi-squared test statistic of 2.7252 and a p-value of 0.4360, we observed no statistically significant difference between the distribution of expected vs observed nuclei counts.

Given that 428 / 765 nuclei were mCherry tagged, this would mean that 28/(428+28)=6.14% percent of true EC-amp were not mCherry tagged. This would therefore set the percentage of true EC-amp nuclei that were thresholded as mCherry tagged at 93.86%.

## Acknowledgements

This work was delivered as part of the eDyNAmiC team supported by the Cancer Grand Challenges partnership funded by Cancer Research UK (CGCATF-2021/100012+ CGCATF-2021/100025) and the National Cancer Institute (OT2CA278688+OT2CA278635). The research was also supported in part by grants U24CA264379, R01GM114362 and Boundless Bio, inc. to VB. X.Y. is a Damon Runyon Fellow supported by the Damon Runyon Cancer Research Foundation (DRG-2474-22). The project described was supported, in part, by Award Number 1S10OD010580-01A1 from the National Center for Research Resources (NCRR). Its contents are solely the responsibility of the authors and do not necessarily represent the official views of the NCRR or the National Institutes of Health. Cell Sciences Imaging Facility provided training and usage support.

## Code Accessibility

InterSeg, interSeg submodules (ecSeg-i and ecSeg-c), and Stat-FISH software are publicly available on GitHub for academic use at https://github.com/UCRajkumar/ecSeg.

