## Supplementary Figures for "Accurate Prediction of ecDNA in Interphase Cancer Cells using Deep Neural Networks"

#### Supplementary Figure Captions

**Supplementary Figure 1: Examples of images from each acquisition protocol.** (a) Image from the cultured cell line GBM39EC (*EGFR* red, Centromere 7 green). (b) Image from the tissue model cell line DLD1 (*MYC* red, Centromere 8 green). (c) Image from a neuroblastoma patient tissue sample (*MCYN* green, Centromere 2 red).

**Supplementary Figure 2: EcSeg-c Architecture Diagram.** EcSeg-c architecture based on DenseNet-121.

**Supplementary Figure 3: Stat-FISH pipeline.** Visualization of stat-FISH pipeline and sample analysis.

**Supplementary Figure 4: EcSeg-i's filters and feature maps.** Top left shows a sample image (*MYCN* red and Centromere 2 green) that is passed to ecSeg-i. Top right shows filters from the first convolutional layer. Bottom row shows the feature maps produced by convolving the filters over the sample image. The ecSeg-i prediction for this nucleus is EC-amp.

**Supplementary Figure 5: InterSeg evaluation on images with shrinking distortion**. The top row shows the original HSR-amp image, the shrunken image, and the interSeg predictions for both. The bottom row shows the original EC-amp image, the shrunken image, and the bootstrapped interSeg predictions for both.

**Supplementary Figure 6: InterSeg evaluation on images with enlarging distortion.** The top row shows the original HSR-amp image, the enlarged image, and the interSeg predictions for both. The bottom row shows the original EC-amp image, the enlarged image, and the bootstrapped interSeg predictions for both.

**Supplementary Figure 7: InterSeg evaluation on images with rotation distortion.** The top row shows the original HSR-amp image, the rotated image, and the interSeg predictions for both. The bottom row shows the original EC-amp image, the rotated image, and the bootstrapped interSeg predictions for both.

**Supplementary Figure 8: InterSeg evaluation on HSR-amp images with contrast distortion.** The top row shows the original HSR-amp image, a low contrast version, and a high contrast version. The middle row displays the pixel distribution of the blue and red color channels to quantify the contrast. The bottom row shows the bootstrapped interSeg predictions, with 'n' representing the number of cells analyzed.

**Supplementary Figure 9: InterSeg evaluation on EC-amp images with contrast distortion.** The top row shows the original EC-amp image, a low contrast version, and a high contrast version. The middle row displays the pixel distribution of the blue and red color channels to quantify the contrast. The bottom row shows the bootstrapped interSeg predictions, with 'n' representing the number of cells analyzed.

**Supplementary Figure 10: InterSeg performance without bootstrapping.** (a) F1-score on the cultured and tissue model test set, where n is the number of cells in each class. (b) Raw (non-bootstrapped) distribution of amplification mechanism of no-amp cell lines. (c) Raw distribution of amplification mechanism of EC-amp cell lines. (d) Raw distribution of amplification mechanism of HSR-amp cell lines.

**Supplementary Figure 11: InterSeg prediction of ecDNA in SF268 interphase cells.** The left image depicts interphase nuclei and metaphase spread. The *MMP8* oncogene is visualized by the red FISH probe, while pan-centromeres are shown by the green FISH probe. In the middle image, a zoomed-in view of a selected nucleus is presented. The right image displays the segmented nucleus, as identified by stat-FISH, showing peaks of the oncogene and centromere signals. Below the right image, we present interSeg predictions for the respective nucleus and stat-FISH results indicating the number of oncogene foci in the nucleus and metaphase.

**Supplementary Figure 12: InterSeg prediction of HSR-amp in SF268 interphase cells.** The left image depicts another metaphase spread of SF268 containing intact nuclei. The *MMP8* oncogene is visualized by the red FISH probe, while pan-centromeres are shown by the green FISH probe. In the middle image, a zoomed-in view of a selected nucleus is presented. The right image displays the segmented nucleus, as identified by stat-FISH, showing peaks of the oncogene and centromere signals. Below the right image, we present interSeg predictions for the respective nucleus and stat-FISH results indicating the number of oncogene foci in the nucleus and metaphase.

**Supplementary Figure 13: InterSeg prediction of ecDNA in SN12C interphase cells.** The left image depicts a metaphase spread of SN12C containing intact nuclei. The *TNFRSF13B* oncogene is visualized by the red FISH probe; the green FISH probe is pan-centromeric. In the middle image, a zoomed-in view of a selected nucleus is presented. The right image displays the segmented nucleus, as identified by stat-FISH, showing peaks of the oncogene and centromere signals. Below the right image, we present interSeg predictions for the respective nucleus and stat-FISH results indicating the number of oncogene foci in the nucleus and metaphase.

**Supplementary Figure 14: Visualization of heterogeneous mix of COLO320DM and COLO320HSR.** The top row of images depicts 3 nuclei in the hybrid COLO320DM and COLO320HSR plate with the mCherry RFP probe in red and the DNA-FISH probe for *MYC* in green. The bottom row displays only the mCherry RFP probe in grayscale, with the color bar displaying observed pixel intensity. Based on the distribution of the green FISH-probe for *MYC*, the nucleus in the left column appears to be HSR-amplified, while the middle and right column nuclei appear to be EC-amplified. While the middle column nucleus displays a high mCherry signal, the right column nucleus displays a low mCherry signal.

**Supplementary Figure 15: InterSeg prediction accuracy in heterogeneous mix of COLO320DM and COLO320HSR.** The left and right boxplots display the precision vs recall curves for mCherry tagged nuclei (COLO320DM) and not mCherry tagged nuclei (COLO320HSR), respectively. Each point represents a threshold for the maximum mCherry brightness per nuclei. Nuclei below the threshold were labeled as not mCherry tagged

(HSR-amp) and the remaining nuclei were labeled as mCherry tagged (EC-amp). For each choice of threshold, the mCherry tagging status was used as a gold-standard to measure the accuracy of interSeg prediction.

**Supplementary Figure 16: InterSeg prediction on neuroblastoma hold-out set images without bootstrapping.** Non-bootstrapped distribution of interSeg amplification mechanism across the 67 NB hold-out test set images, stratified by the pathologist annotated ‘amplification’ and 'No amplification' labels. Each column corresponds to a single patient, and the bar height corresponds to the proportion of cells labeled for each amplification class by interSeg.

### Supplementary Figure 1

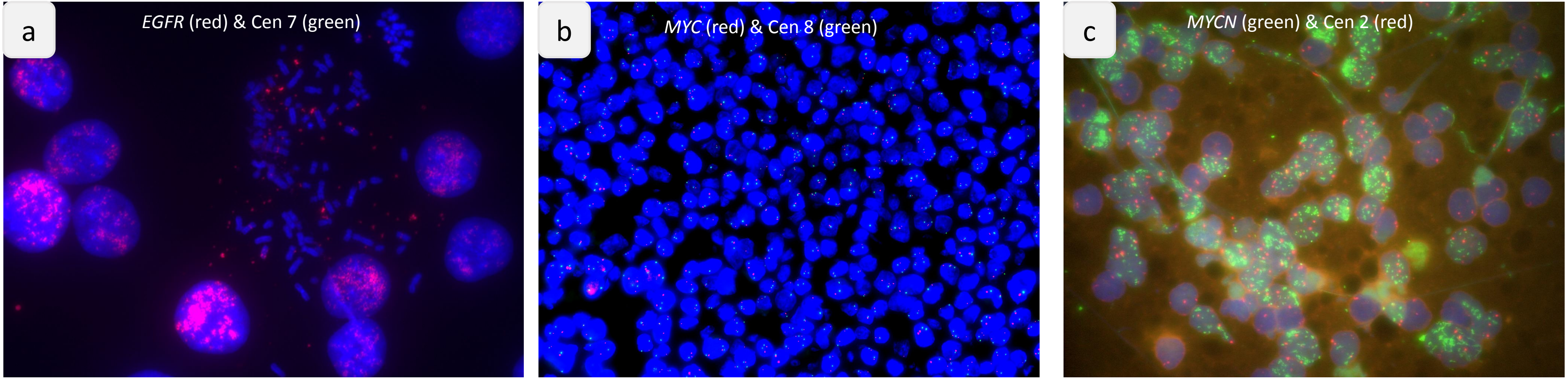


Cultured cells Tissue model Patient tissue

### Supplementary Figure 2

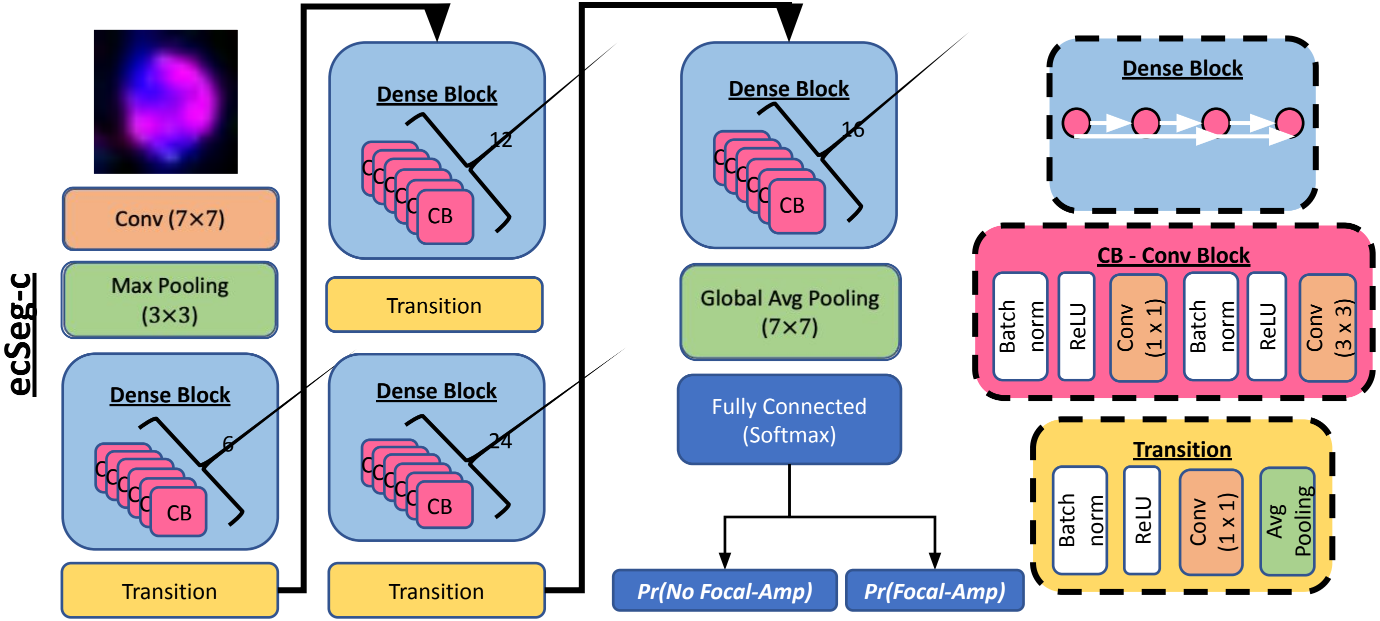

### Supplementary Figure 3

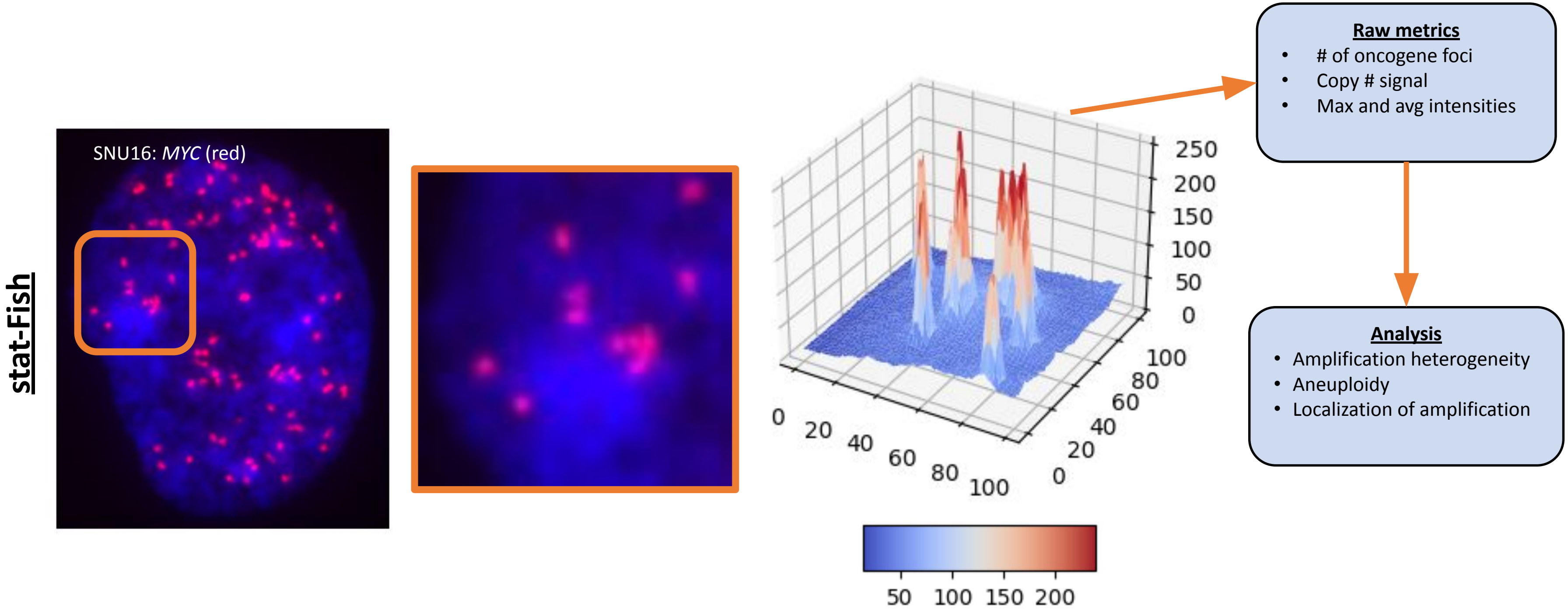

### Supplementary Figure 4

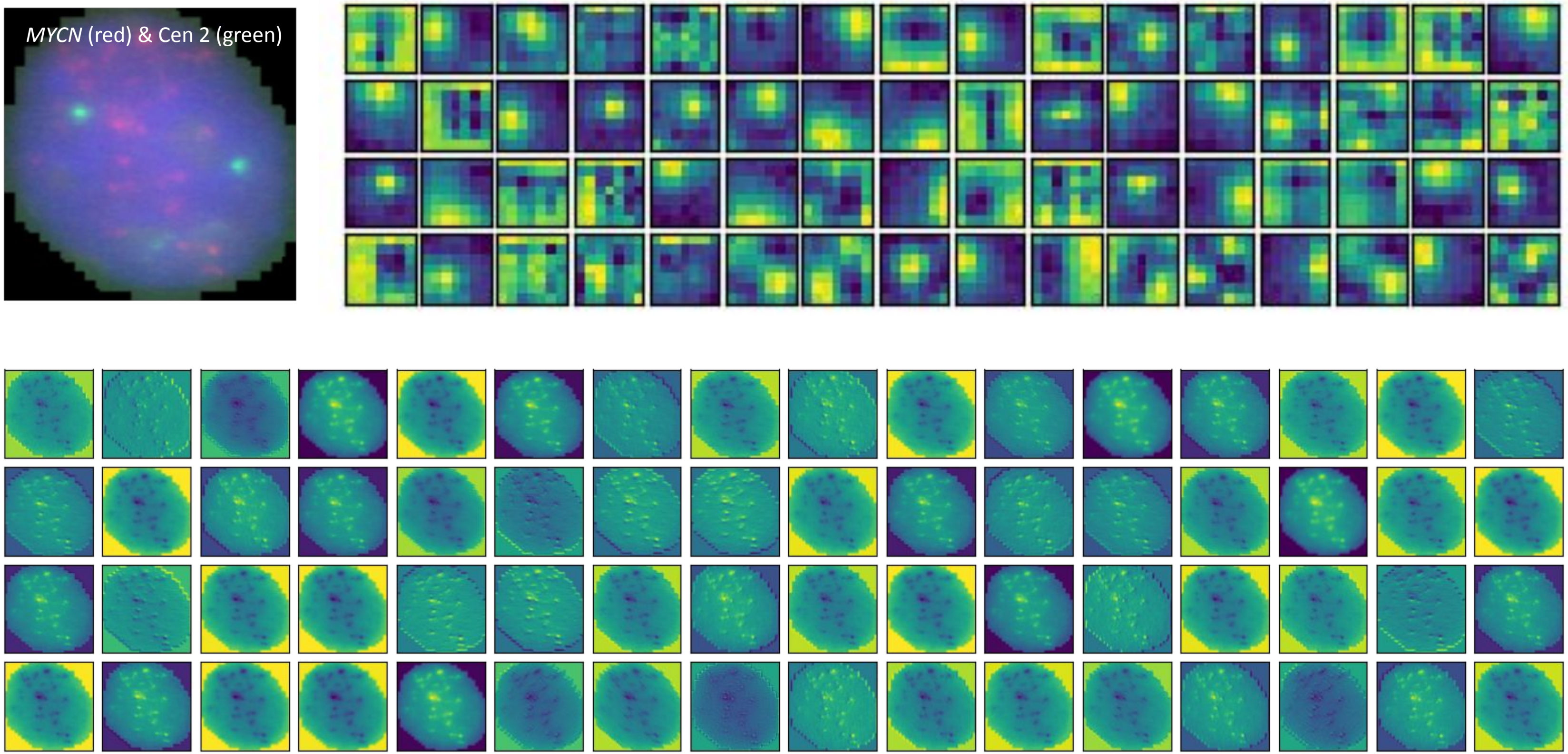

### Supplementary Figure 5

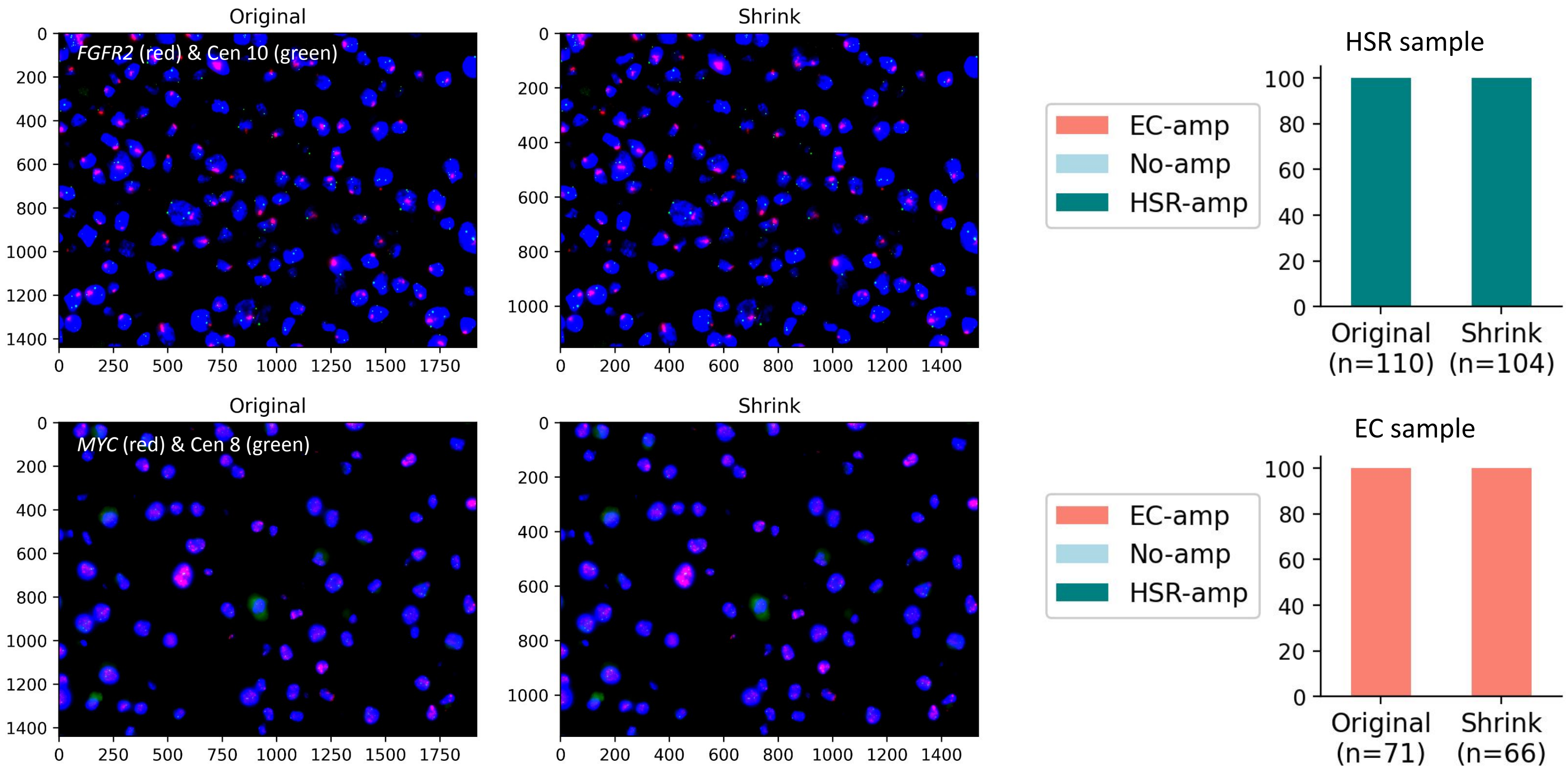

### Supplementary Figure 6

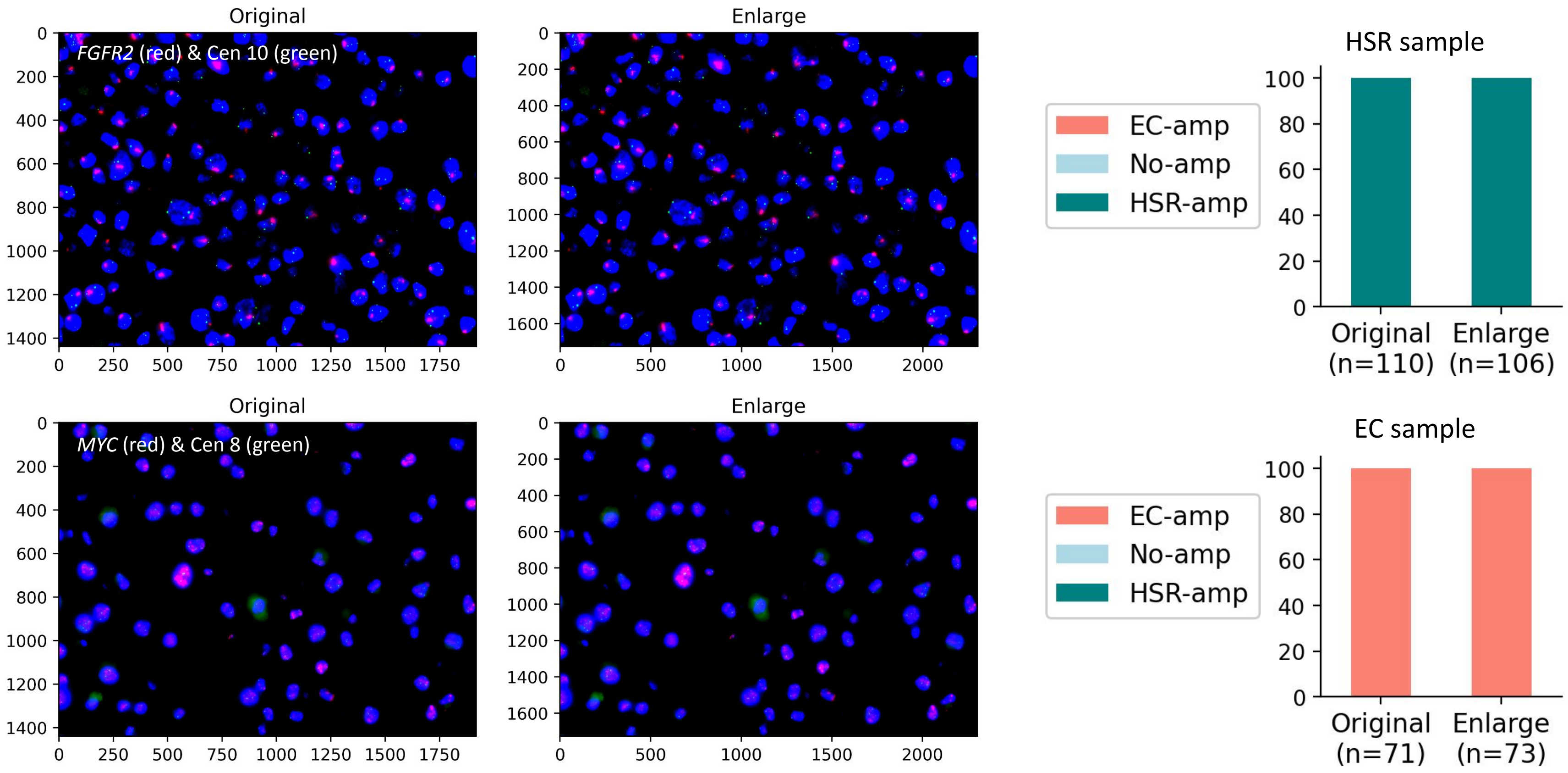

### Supplementary Figure 7

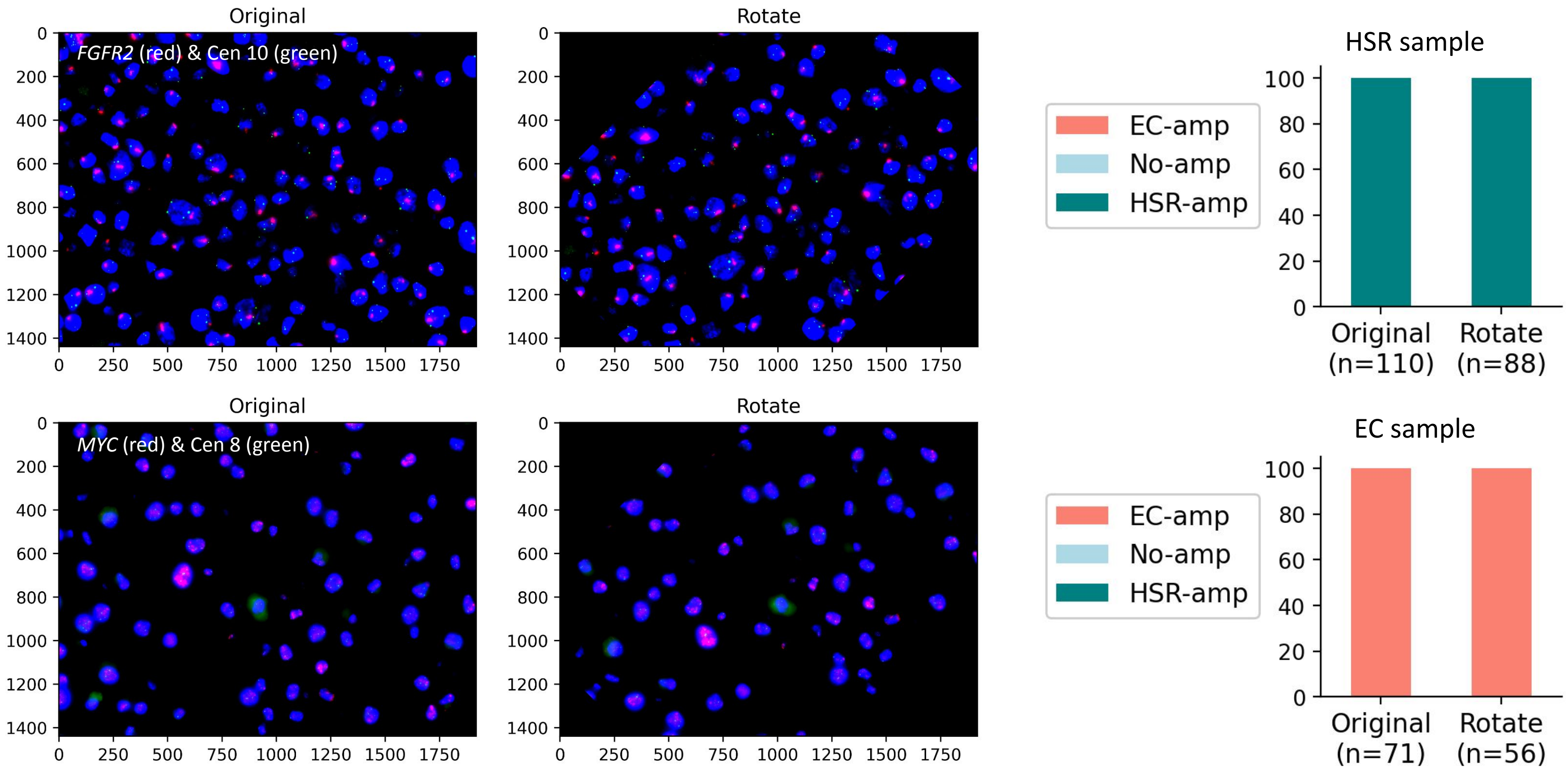

### Supplementary Figure 8

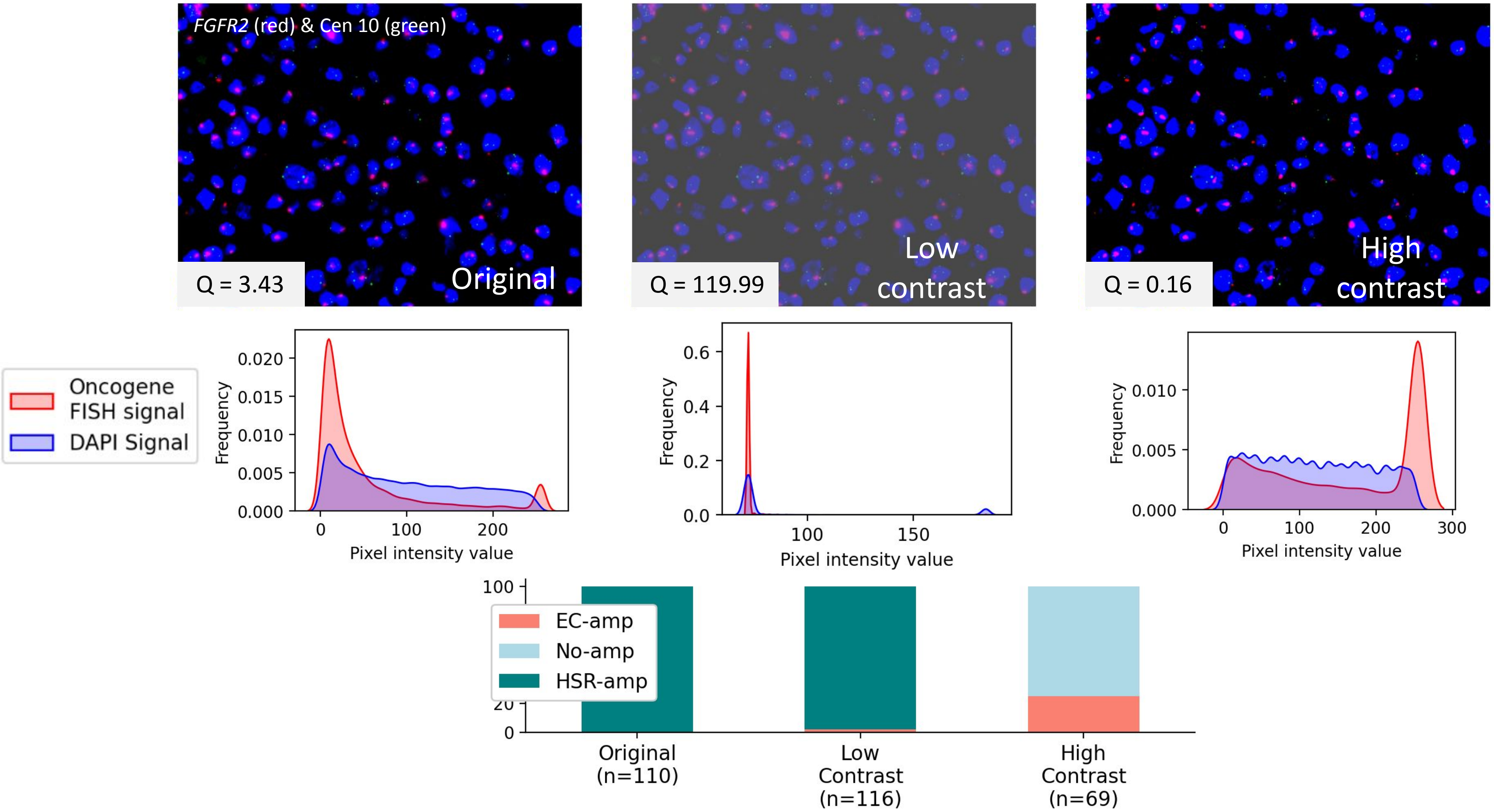

### Supplementary Figure 9

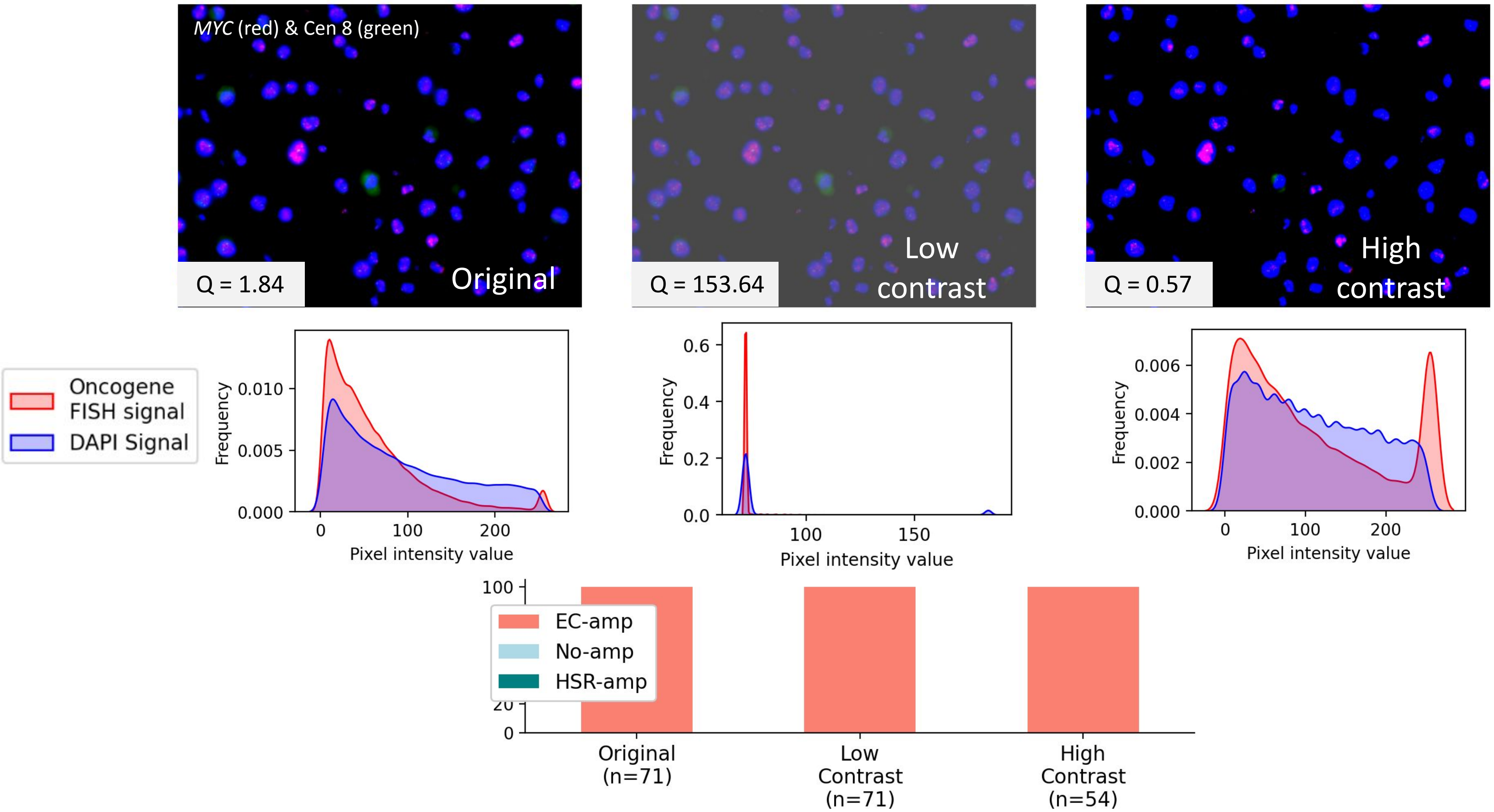

### Supplementary Figure 10

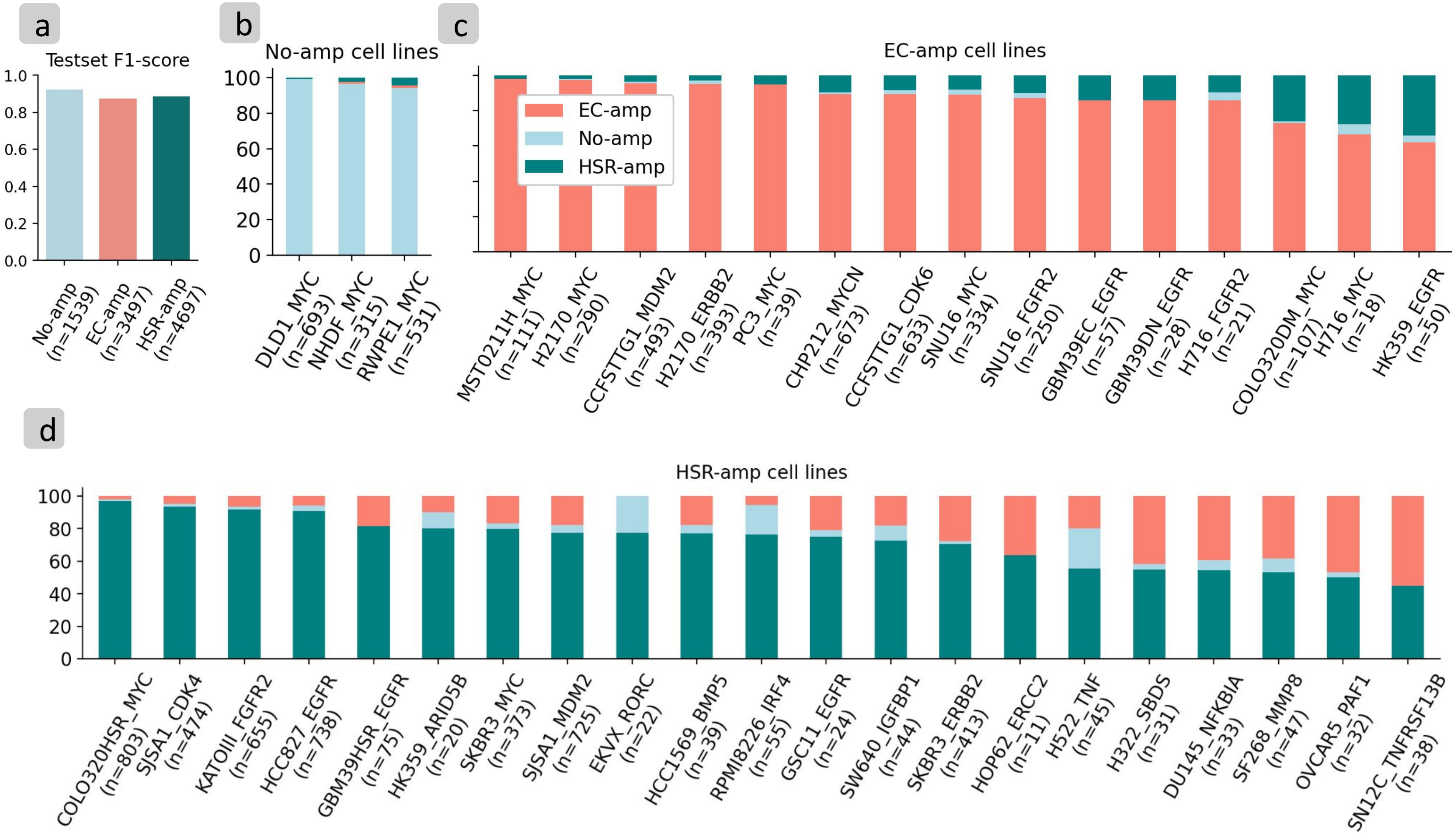

### Supplementary Figure 11

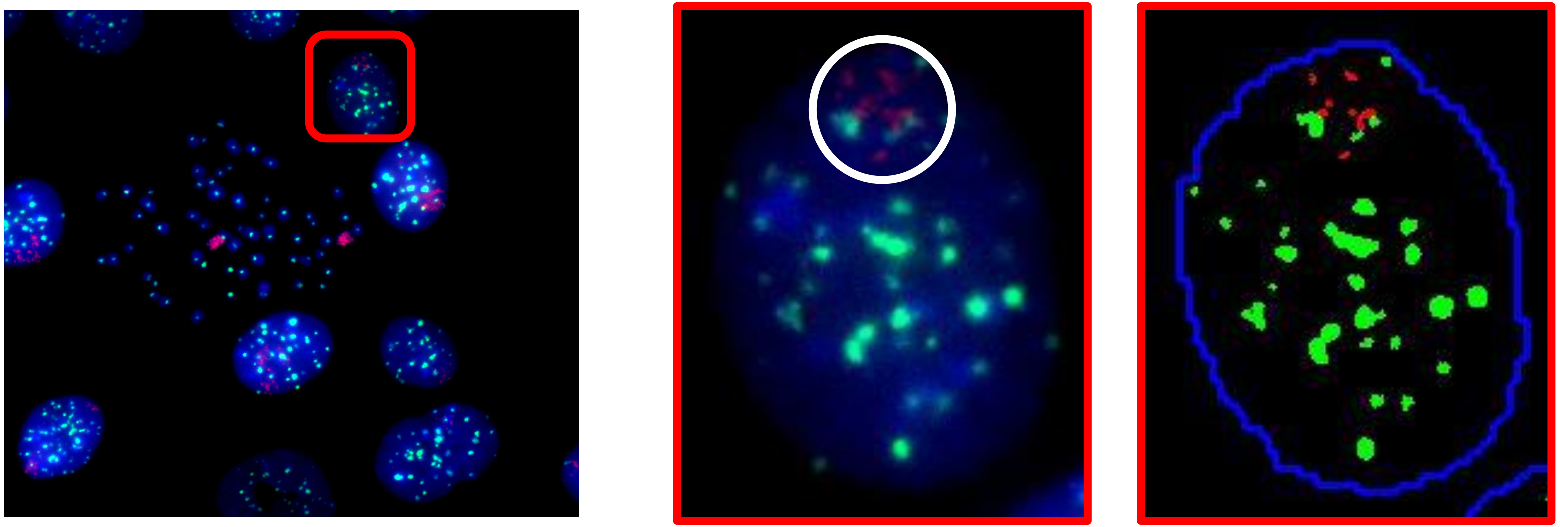

Cell line: SF268
Pan-centromeric probe (green)
*MMP8* oncogene (red)

P(No-amp) = 3.2e-08
P(EC-amp) ≈ 1.0
P(HSR-amp) = 2.4e-07

Number of foci in metaphase: 2
Number of foci in interphase: 7

### Supplementary Figure 12

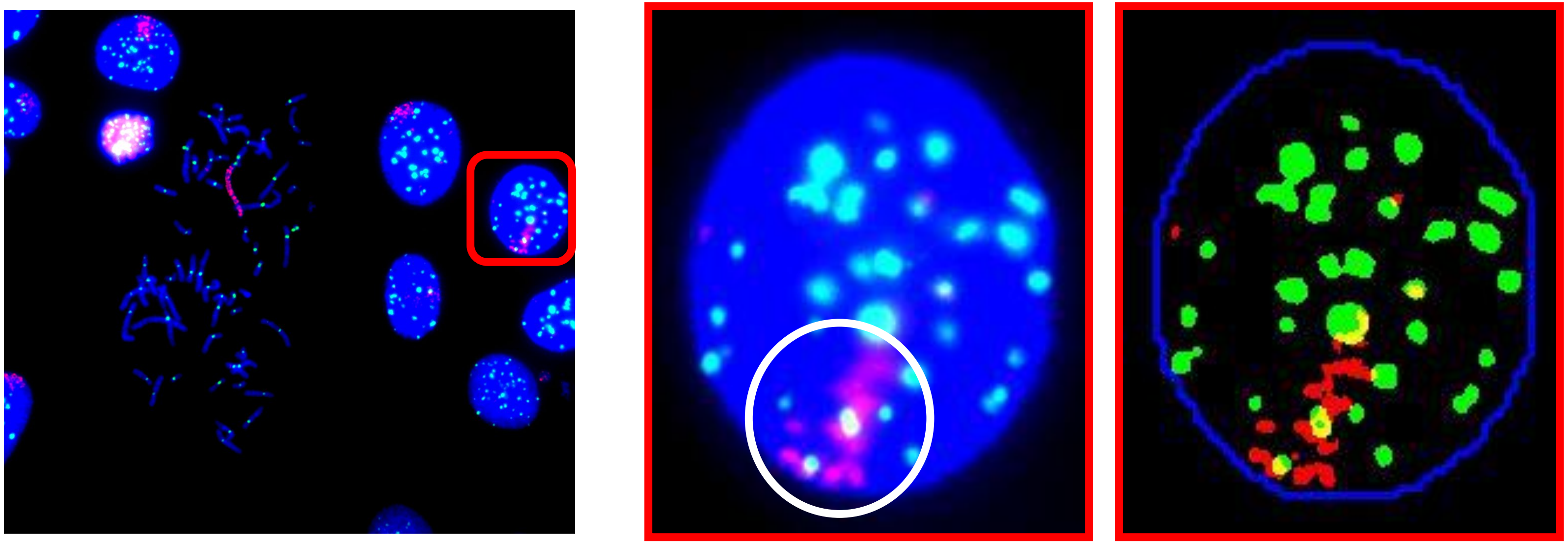

Cell line: SF268
Pan-centromeric probe (green)
*MMP8* oncogene (red)

P(No-amp) = 1.9e-07
P(EC-amp) = 1.0e-3
P(HSR-amp) ≈ 1.0

Number of foci in metaphase: 2
Number of foci in interphase: 9

### Supplementary Figure 13

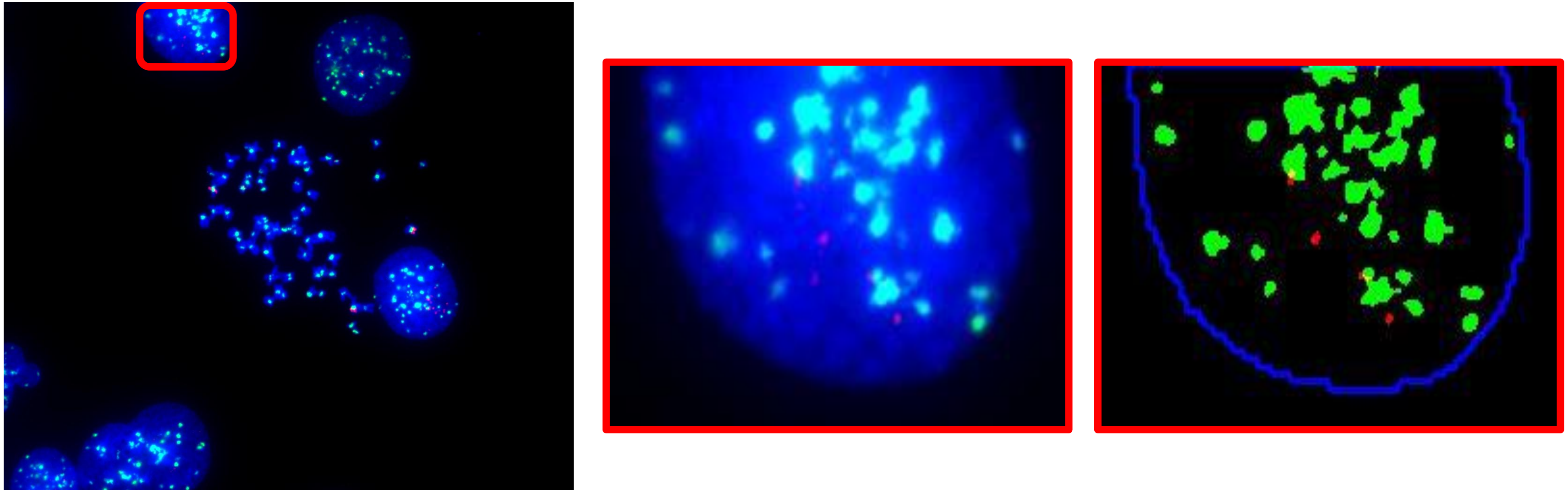

Cell line: SN12C
Pan-centromeric probe (green)
*TNFRSF13B* oncogene (red)

P(No-amp) = 8.5e-13
P(EC-amp) ≈ 1.0
P(HSR-amp) = 2.7e-09

Number of foci in metaphase: 4
Number of foci in interphase: 4

### Supplementary Figure 14

Likely HSR-amp with low mCherry | Likely EC-amp with high mCherry | Likely EC-amp with low mCherry

COLO320DM/HSR
*MYC* (green)
mCherry (red)

COLO320DM/HSR
mCherry Channel

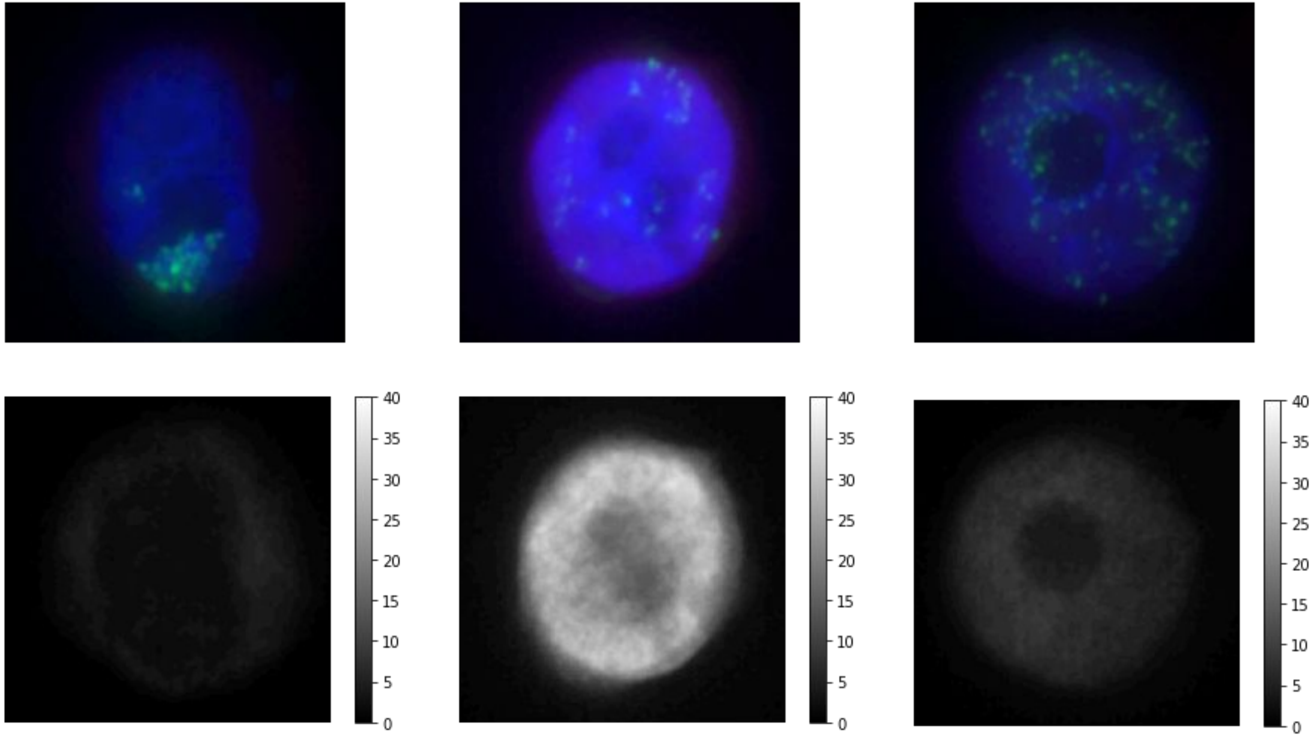

### Supplementary Figure 15

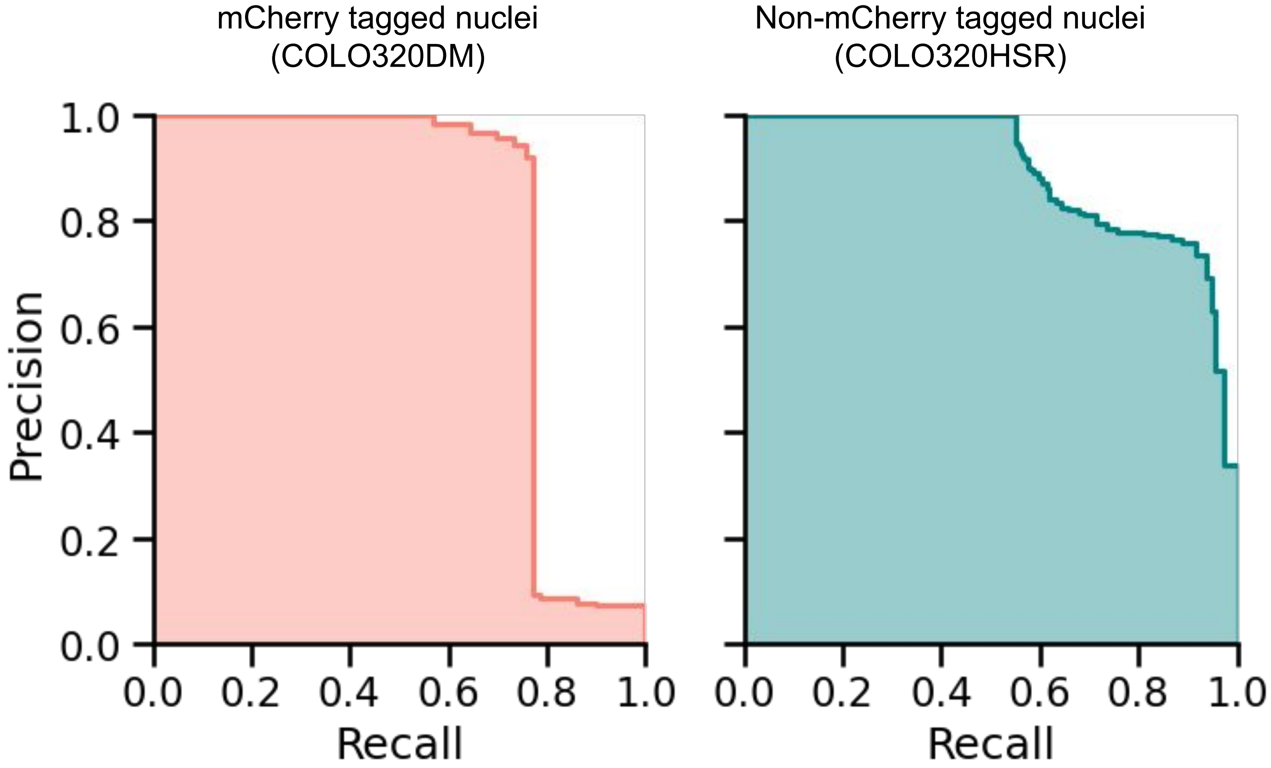

### Supplementary Figure 16

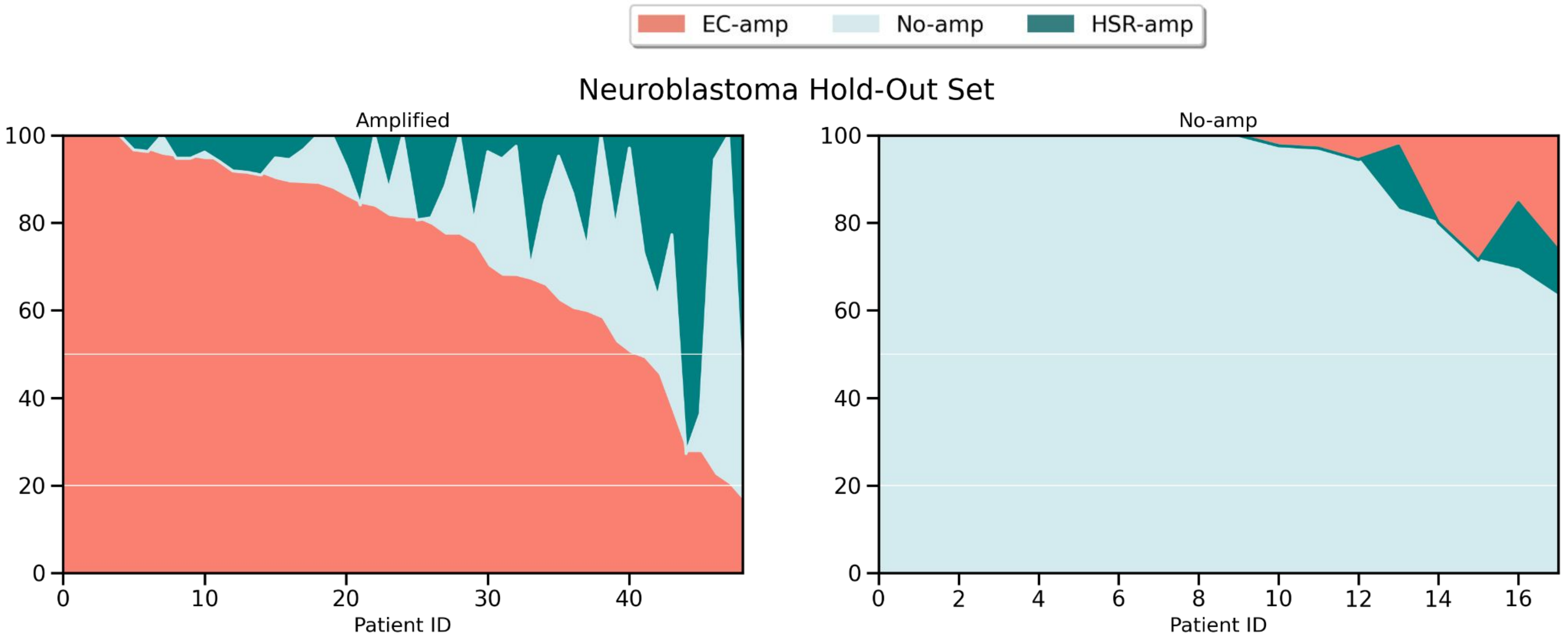
